# scTour: a deep learning architecture for robust inference and accurate prediction of cellular dynamics

**DOI:** 10.1101/2022.04.17.488600

**Authors:** Qian Li

## Abstract

Despite the continued efforts to computationally dissect developmental processes using single-cell genomics, a batch-unaffected tool that is able to both infer and predict the underlying dynamics is lacking. Here, I present scTour, a novel deep learning architecture to perform robust inference and accurate prediction of the cellular dynamics in diverse processes. For inference, scTour can efficiently and simultaneously estimate the developmental pseudotime, intronic read-independent vector field, and transcriptomic latent space under a single, integrated framework. For prediction, scTour can precisely reconstruct the underlying dynamics of unseen cellular states or an independent dataset agnostic to the model. Of note, both the inference and prediction are invariant to batch effects. scTour’s functionalities are successfully applied to a variety of biological processes from 17 datasets such as cell differentiation, reprogramming and zonation, providing a comprehensive infrastructure to investigate the cellular mechanisms underpinning development in an efficient manner.

## Introduction

Amongst the challenges that decoding developmental processes at single-cell resolution using single-cell RNA sequencing (scRNA-seq) poses, a unique difficulty is that scRNA-seq can only capture static snapshots of cells. In addition, experimental assays such as lineage tracing and metabolic labelling are inaccessible to many biological systems particularly those involving human tissues^1–5^. Many computational tools have been developed to analyse these dynamic processes, the most prevalent of which are pseudotime-based ordering of cells along their trajectory and RNA velocity-based directing of future cell states^6–10^. Despite the wide usefulness of these tools, they have several limitations which restrict their scope: (1) the majority of tools for pseudotime estimation require the users to explicitly designate the starting cells, meaning that they are limited to well-studied biological processes. (2) the existing RNA velocity-based tools are largely focused on the modelling of transcriptional kinetics. This requires either extraction of spliced and unspliced mRNAs within cells, a rate-limiting step especially for large-scale datasets, or information from metabolic labelling which is often not possible especially when applied to human tissues^9^. This could also lead to inaccurate inference due to the assumption of constant kinetic rates and the noisy approximation of nascent transcripts by intronic reads^11^. Moreover, they are not readily adaptable to use cases beyond scRNA-seq. (3) current algorithms are affected by batch effects to varying degrees and thus demand batch corrections prior to the formal analyses. This is particularly difficult for time-course experiments. (4) the prediction functionality is lacking or quite limited in the current methods. Neither the pseudotime nor the vector field can be made predictable for unseen data. Although two recent studies did use the vector field to predict the transcriptomic space forward or backward given an initial cell state^9, 12^, predicting unseen cellular states is challenging for these tools. All these issues restrict the current methods to the data they have modelled and hinder the transfer and generalization to new datasets.

Here I introduce scTour, an innovative deep learning-based architecture that, in addition to overcoming the limitations detailed above, achieves multifaceted dissection of a variety of biological processes under a single model. scTour simultaneously infers the developmental pseudotime, transcriptomic vector field and latent space of cells, with all these inferences unaffected by batch effects inherent in the datasets. Another advantage is that the pseudotime estimation does not require the indication of a starting cell, and the vector field inference does not rely on the discrimination between spliced and unspliced mRNAs, rendering scTour applicable to other genomic data. Importantly, the inference of a low-dimensional latent space which combines the intrinsic transcriptome and extrinsic time information provides richer information for reconstructing a finer cell trajectory. Uniquely in scTour, the resulting model can be further employed to predict the transcriptomic properties and dynamics of unseen cellular states and even to predict the characteristics of a different dataset new to the model. These together make scTour a generative and powerful method for single-cell developmental data analysis. To demonstrate the superiority of scTour, I have applied it to a wide variety of dynamic biological processes including neurogenesis, pancreatic endocrinogenesis, skeletal muscle, thymic epithelial cell and embryonic development, hematopoiesis, and brain vasculature zonation (scRNA-seq), as well as reprogramming (single-nucleus RNA-sequencing (snRNA-seq)) and human fetal retinal development (single-cell ATAC-sequencing (scATAC-seq)). In all of these systems, the accuracy and effectiveness of scTour in recapitulating the underlying cellular dynamics was validated. scTour is available as an open-source software at https://github.com/LiQian-XC/sctour.

## Results

### The scTour architecture

scTour is a new deep learning architecture that builds on the framework of variational autoencoder (VAE)^13^ and neural ordinary differential equation (ODE)^14^ accompanied by critical innovations tailored to the analysis of dynamic processes using single-cell genomic data (Fig. 1). Specifically, given a gene expression matrix, scTour leverages a neural network to assign a time point to each cell in parallel to the neural network for latent variable parameterization. The resulting time information allows scTour to spot the initial latent state *z_t_*_0_, which is further combined with the estimated time of each cell to solve an ODE, with the derivative of latent states with respect to time defined by another neural network (Fig. 1). The ODE solver yields another series of latent representations, together with the one from variational inference, to serve as the input for reconstructing the transcriptomes in a weighted manner (see Methods).

**Fig. 1.**
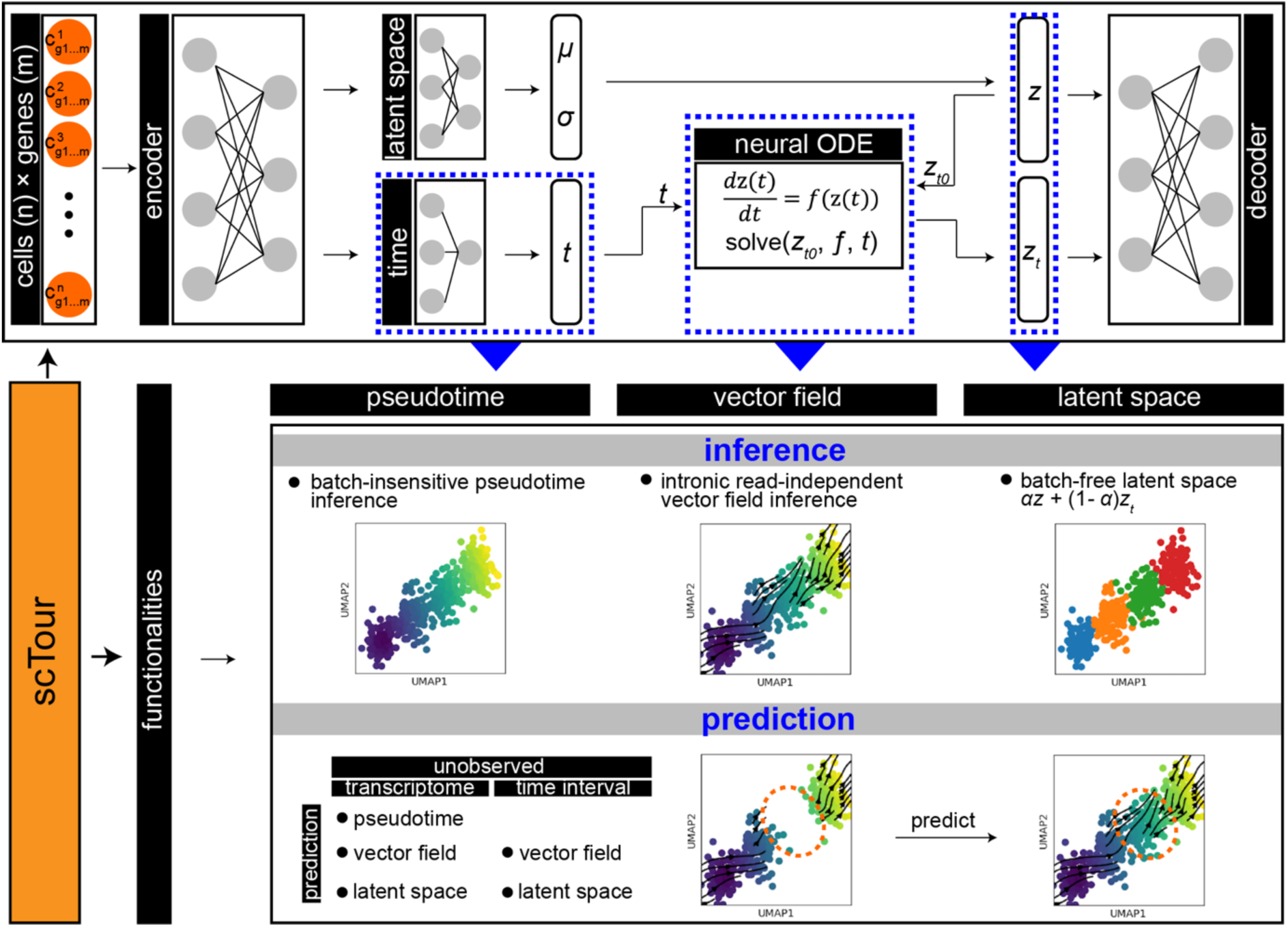
scTour framework. With a gene expression matrix as input, an encoder network is used to both generate the distribution parameters of the approximate posterior (latent space, *z*) and assign a time point to each cell (time, *t*). The sample from the posterior at the initial state (*z_t_*_0_) along with the times (*t*_0_, *t*_1,_ *t*_2_,…,*t_n_*) of cells are input into a neural ODE to yield another series of latent representations *z_t_*. A decoder network then reconstructs the input using the latent *z* and *z_t_*. This model can be used to infer the developmental pseudotime, transcriptomic vector field and latent representations of cells in an unsupervised manner, as well as to predict the cellular dynamics of unobserved transcriptomes or time intervals.

Compared to the latent ODE model proposed in the original neural ODE publication^14^, scTour delivers three major innovations. Firstly, scTour introduces a neural net for inferring the developmental time of a given cell based on its transcriptome. This operation enables the model to bypass the dependence on the prior knowledge of the cell timeline, and endows scTour with the ability to suit any data beyond the timestamped ones. Secondly, different from the original model, which adopts a recurrent neural network (RNN) as the recognition net to derive the latent state only at time *t*_0_, scTour employs the typical encoder to infer the latent states covering all observations. These are then used to reconstruct the transcriptome space concurrently with the ones from the ODE solver. Such an operation preserves the intrinsic transcriptomic structure of cells and proves a superior strategy in reconstructing the trajectory. Thirdly, scTour utilises the standard mini-batch training which is less straightforward in the original latent ODE model^14^. With this optimization, scTour’s performance is again improved, being highly efficient and scalable to large-scale datasets.

As a result, scTour provides two main functionalities in deciphering cellular dynamics in a batch-unaffected manner: inference and prediction (Fig. 1). For inference, the time neural net in scTour allows estimates of cell-level pseudotime along the trajectory with no need for specifying starting cells. The learned differential equation (i.e., the latent state’s derivative with respect to time) by another neural net provides an alternative way of inferring the transcriptomic vector field. This eliminates the time-consuming step of distinguishing spliced from unspliced mRNAs used in RNA velocity-based tools and thus can be extended to other genomic data. The variational inference and ODE solver yield a combined latent representation which contains richer information for both reconstructions of developmental trajectories and cellular stratifications. For prediction, given an unobserved cellular state or a new dataset agnostic to the model, the time neural net trained in scTour can predict its developmental pseudotime; the learned differential equation can infer its transcriptomic vector field; the latent space is likewise predictable. Notably, the latent space of an unseen cellular state can also be reconstructed by providing the model with its expected developmental time. All these are novel and powerful features adding to the existing trajectory inference tools.

### scTour’s inference captures the underlying developmental dynamics

I first evaluated scTour using a scRNA-seq dataset from the mouse dentate gyrus during postnatal development. The focus here was on the granule cell lineage which undergoes sequential transcriptomic changes from neuronal intermediate progenitor cells (nIPCs), neuroblasts, immature granule cells, to mature granule cells^15^ (4,007 cells, Fig. 2a). Following the scTour model training (see Methods), the developmental pseudotime, transcriptomic vector field, and low-dimensional latent space (set as five dimensions) of cells were derived (Fig. 2a). The estimated pseudotime clearly recapitulated the developmental process of granule cells, with the transcriptional continuum from nIPCs to mature granule cells captured. Similarly, analysis of the vector field delineated the expected directional flow along the differentiation path when visualised on the uniform manifold approximation and projection (UMAP) embedding (Fig. 2a). Thus, scTour consists of two complementary approaches to dissect the course of cell differentiation. An important advantage is that the vector field, which was solely derived from the expression matrix without the need for investigation of spliced and unspliced transcripts, assessed the cellular dynamics more efficiently. At the same time it performed better than the intronic read-based velocity estimate which failed to capture the immature to mature granule cell transition (Supplementary Fig. 1). The latent space computed by scTour through incorporating both the intrinsic transcriptome and extrinsic pseudotime information not only reflected the transcriptomic differences among cell types, but also charted a finer continuous trajectory underlying the developmental process of granule cells when compared to that constructed from the PCA space (Fig. 2a).

**Fig. 2.**
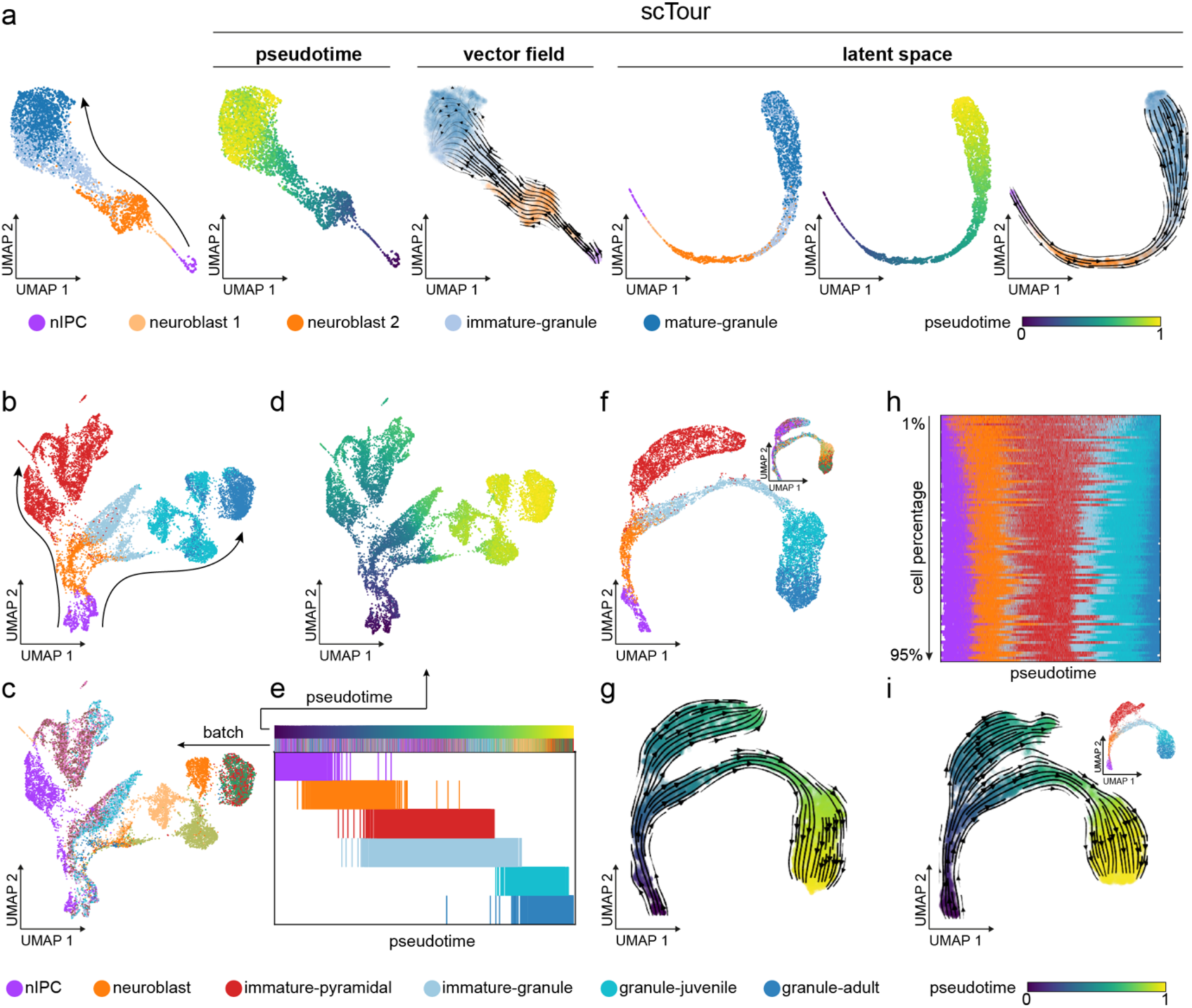
scTour robustly captures the cellular dynamics during dentate gyrus neurogenesis. **a,** UMAP visualizations of cell types from the granule cell lineage (4,007 cells)^15^, and the developmental pseudotime, transcriptomic vector field and latent representations inferred by scTour. Leftmost panel shows the PCA space-based UMAP with the arrow indicating the differentiation from nIPCs to mature granule cells. **b,** PCA space-based UMAP embedding showing cell types (colours, 15,174 cells)^15^ along the pyramidal and granule cell lineages (arrows). **c,** As in **b**, but coloured by sample batches. **d,** As in **b**, but coloured by the developmental pseudotime derived from the scTour model. **e,** Developmental ordering of cells by the pseudotime inferred from scTour. Cells are coloured from top to bottom by pseudotime, sample batches and cell types. **f,** UMAP visualizations of the latent representations learnt from scTour, with colours denoting the cell types and sample batches (top right). **g,** Streamline visualization of the transcriptomic vector field from scTour on the same embedding as in **f**, with cells colour-coded by the inferred pseudotime. **h,** Developmental ordering of cells by the pseudotime estimated from scTour models trained using a range of cell subsets (1% to 95% of total cells from top to bottom). Cells are coloured by cell types. **i,** UMAP visualizations of the latent representations, developmental pseudotime (colours), and transcriptomic vector field (streamlines) learnt from the scTour model trained based on 20% of total cells. The top- right panel shows the same plot colour-coded by cell types.

### scTour’s inference is invariant to batch effects and cell subsampling

The advantages of applying scTour to a linear and continuous developmental process are clear. To further test its capability in dealing with more complex processes, I next applied scTour to another scRNA-seq dataset from the developing mouse dentate gyrus which collected some extra immature pyramidal neurons from the hippocampus proper^15^. I focused on the granule cell lineage along with the immature pyramidal neurons; in the original study it was suggested that they shared a differentiation trajectory (15,174 cells, Fig. 2b). This dataset presented substantial batch effects from different samples that segregated cells significantly within the same cell type (Fig. 2c). Nevertheless, scTour successfully recapitulated the two differentiation branches without any impact from the sample batches due to the continuous-in-time transformations of the latent states by the ODE solver (Fig. 2d-g). Specifically, the estimated pseudotime was in line with the differentiation courses, depicting the gradual progression from nIPCs to both granule cells and pyramidal neurons (Fig. 2d,e). The inferred latent space was also batch free and constructed an improved cell differentiation trajectory (Fig. 2f). Projecting the vector field onto this trajectory further, scTour again corroborated the shared trajectory between granule and pyramidal cell lineages, with the immature parts of both cell populations branching out from the neuroblasts (Fig. 2g). This feature of scTour is of critical importance to cross-platform or cross-study data integrations and comparisons because it is not conditioned on batch corrections and thus alleviates the risk of overcorrection when batch confounders and biological signals are entangled (such as two organs from two individuals respectively). Altogether, scTour’s inference of cellular dynamics is batch insensitive, and thus provides an easy and accurate way of interrogating single-cell datasets from multiple angles.

Given scTour’s design of model-based prediction and implementation of mini-batch training, it was possible that a scTour model could be trained from a subset of data and the resulting model could be used to derive the characteristics of the entire dataset. To test this possibility, I trained scTour models on the same dataset but used a series of subsets ranging from 1% to 95% of all cells. The results highlighted the robustness of scTour, as both the granule and pyramidal cell lineages already manifested when the model was from as small as 1% of all cells (Supplementary Fig. 2a-c). Across the subsampling span from 1% to 95%, the inferred full spectrum of cellular transcriptomic dynamics converged quickly (Fig. 2h and Supplementary Fig. 2d). To illustrate this, it was clear that the pseudotime, vector field and latent space learnt from 20% of data successfully reconstructed the full granule and pyramidal cell differentiation paths (Fig. 2i). For all these analyses, since the scTour model was trained with a small subset of cells (20%), it took 12 minutes for the model training using CPU only and one second to propagate to full data inference (15,174 cells). All these endow scTour with remarkable efficiency and scalability when dealing with large-scale datasets.

Taken together, scTour can characterise dynamic processes comprehensively, robustly and efficiently, allowing for its application to diverse datasets from different biological processes, systems, species, and experimental platforms. These include, but are not limited to, mouse embryonic organoids^16^ (30,496 cells, Supplementary Fig. 3), human thymic epithelial cell development^17^ (14,217 cells, Supplementary Fig. 4), human embryonic development^18, 19^ (1,195 cells, Supplementary Fig. 5; 90 cells, Supplementary Fig. 6), induced pluripotent stem cell (iPSC) reprogramming^20, 21^ (251,203 cells, Supplementary Fig. 7; 36,597 nuclei, Supplementary Fig. 8), hematopoiesis^9^ (1,947 cells, Supplementary Fig. 9), and brain vasculature zonation^22^ (3,105 cells, Supplementary Fig. 10). All these analyses demonstrated the efficiency and accuracy of scTour’s inference. A particular advantage of scTour is that the transcriptomic vector field can be directly obtained from single-nucleus data to elucidate the reprogramming process (Supplementary Fig. 8). This is challenging for RNA velocity-based tools due to the disruption of the balance between spliced and unspliced transcripts during the nucleus isolation^11^. Another striking example was the delineation of a dataset focussed on hematopoiesis where the underlying cell trajectory was not captured by the spliced RNA velocity but only by the total RNA velocity from metabolic labelling^9^. With scTour, this process was easily depicted with no dependence on extra information or experimental assays (Supplementary Fig. 9).

### scTour’s prediction reconstructs the dynamics of unseen cellular states

Given the predictive functionality built in scTour, I next assessed its ability to predict the characteristics of unseen cellular states (i.e., cellular states new to the model). I therefore applied scTour to a scRNA-seq dataset from the development of endocrine compartment of the mouse pancreas, as previously described in the scVelo publication^8, 23^ (3,696 cells). The mouse pancreatic endocrinogenesis starts from the endocrine progenitors (EPs), goes through the intermediate stage (*Fev*+ endocrine cells), and finally commits to four major fates: α-cells, β- cells, δ-cells, and ε-cells. I started by training the scTour model using all the cellular states involved in this process. Here I compared the derived developmental pseudotime with scVelo’s latent time. This was because the latter was shown to delineate this process more accurately than diffusion pseudotime as it captured the earlier emergence of α-cells relative to β-cells^8^. This comparison highlighted the usefulness of scTour’s pseudotime in not only resolving the ordering of α- and β-cells, but also identifying the continuous progression from *Fev*+ endocrine cells to terminal fates which was not revealed by scVelo’s latent time (Fig. 3a and Supplementary Fig. 11a,b).

**Fig. 3.**
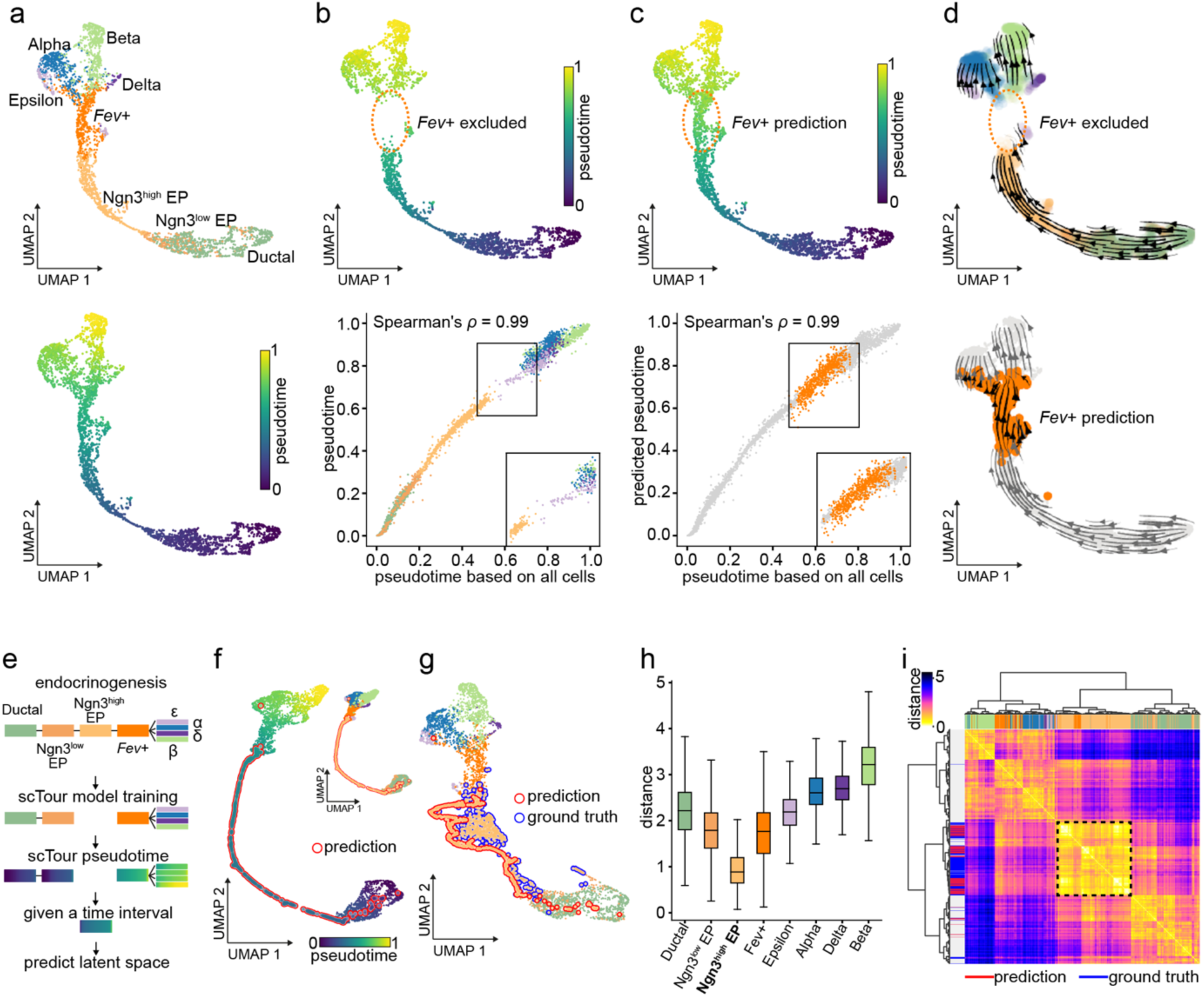
scTour reconstructs the cellular dynamics of unseen cellular states in pancreatic endocrinogenesis. **a,** UMAP visualizations of the latent space inferred by the scTour model which is trained based on 3,696 cells from the endocrinogenesis process^23^. Cells are coloured by cell types (upper) and developmental pseudotime (bottom) estimated from the same model. **b,** Upper panel: UMAP representation showing the pseudotime estimated from the scTour model trained with the *Fev*+ endocrine cells excluded. Lower panel: scatter plot comparing the pseudotime estimate (y axis) with those inferred from the full dataset (x axis). The Spearman correlation coefficient between the two estimates is shown on top and the rectangles mark the gap corresponding to the excluded *Fev*+ endocrine cells. **c,** Upper panel: UMAP representation displaying the predicted pseudotime for the held-out *Fev*+ endocrine cells (dotted circle). Lower panel: scatter plot showing the comparison of the prediction (y axis) with the ground truth (x axis) highlighted in orange. The Spearman correlation coefficient between the two sets of pseudotime in x and y axes is shown on top and the rectangles mark the gap filled by the scTour prediction. **d,** Streamline visualizations of the transcriptomic vector field from the model trained by excluding *Fev*+ endocrine cells (upper), and the predicted vector field for these held-out cells (bottom). **e,** Schematic depicting the scTour model training with the Ngn3^high^ EPs excluded, followed by prediction of the latent space of these cells given their expected developmental time. **f,** UMAP visualization based on the reconstructed latent representations for the held-out Ngn3^high^ EPs (red outline) and those inferred from training cells. Cells are coloured by their developmental pseudotime and cell type identities (top right). **g,** UMAP visualization based on the latent representations of reconstructed Ngn3^high^ EPs (red outline), true Ngn3^high^ EPs (blue outline), and the remaining cells. Cells are coloured by cell types. **h,** Box plot displaying the Euclidean distances calculated between the reconstructed latent representations for Ngn3^high^ EPs and those from each of the cellular states (i.e., true Ngn3^high^ EPs and the remaining states), with the medians, interquantile ranges, and 5th, 95th percentiles indicated by centre lines, hinges, and whiskers, respectively. **i,** Unsupervised hierarchical clustering of the reconstructed Ngn3^high^ EPs along with all the other cells based on their Euclidean distances in the scTour latent space. Column colours of the heatmap mark the cell types and row colours denote the reconstructed (red), true (blue) Ngn3^high^ EPs, and remaining cells (light grey). The colour gradient of the heatmap indicates the Euclidean distance.

Next, I excluded one cellular state, the intermediate *Fev*+ endocrine cells, and trained a scTour model on the remaining cells. The aim was to test: (1) whether scTour can infer the cellular dynamics of a discontinued process; and (2) whether the resulting model can be used to predict the properties of the held-out cellular state. This analysis demonstrated that scTour can recapitulate the discontinuous differentiation course, assigning near-identical pseudotime as compared to that from the analysis of the entire dataset (Fig. 3b), as well as presenting a time gap between EPs and the four terminal states as expected (Fig. 3b). By contrast, scVelo’s latent time was unable to delineate this discontinuous process in full as it failed to disentangle the continuum of early progenitor cells and to recognize the intermediate transitional process by erroneously connecting EPs with terminal states (Supplementary Fig. 11c,d).

On the basis of the model trained above, scTour successfully predicted pseudotime of the unseen cellular state - in this case the *Fev*+ endocrine cells - filling in the time gap and thus bridging the EPs and terminal cells (Fig. 3c). In parallel, the predicted transcriptomic vector field for this cell type correctly orientated those cells towards terminal fates (Fig. 3d). Moreover, scTour predicted the latent space of those unseen cells and, based on this, reconstructed the full trajectory of endocrinogenesis by placing them properly along the differentiation path (Supplementary Fig. 12). In addition to the intermediate cellular states, scTour was capable of reconstructing the dynamics of unobserved starting or terminal states (Supplementary Fig. 12). Taken together, scTour can perform precise out-of-distribution predictions beyond the inference.

### scTour reconstructs the transcriptomic space at unobserved time intervals

During development some intermediate cell states are often transient or present in small quantities. Reconstructing transcriptomic signatures of these cells will be useful when there is limited coverage of particular cell types. scTour allows inference of the transcriptomic characteristics of uncaptured cellular states based merely on their expected developmental time, achieved by integrating the ODE in a stepwise manner and taking into account the *k*- nearest neighbours in the time space when inferring the latent representation at an unobserved time point (see Methods). To test this functionality, a scTour model was trained using the same dataset of pancreatic endocrinogenesis described above but with Ngn3^high^ EPs located between Ngn3^low^ EPs and intermediate *Fev*+ endocrine cells excluded. After training, scTour correctly assigned the developmental pseudotime to each cell, leaving an anticipated time gap corresponding to the missing Ngn3^high^ EP population (Fig. 3e).

Next, when this time interval was provided as the only input to the trained scTour model, the transcriptomic latent space corresponding to this time span was reconstructed and shown to locate at the expected position between Ngn3^low^ EPs and *Fev*+ endocrine cells, forming a complete continuous trajectory together with other cells (Fig. 3f). Of note, this was a rather long-range prediction covering an entire cellular state. When further projecting all the cells onto the same UMAP embedding, the reconstructed and ground-truth Ngn3^high^ EPs were placed together, indicating their transcriptomic similarity (Fig. 3g). This was reinforced by their shortest distance through the comparison with each cellular state in the latent space, revealing the expected trend of transcriptomic difference following the differentiation progression (Fig. 3h). More specifically, unsupervised clustering using the derived distances rebuilt a tree which not only revealed the developmental relations among cell types but also grouped the predicted and true Ngn3^high^ EPs into a single branch (Fig. 3i). All these results illustrated the accuracy of scTour in reconstructing the transcriptomic space at unobserved intermediate time intervals. Besides, scTour can be leveraged to recover the unobserved starting and terminal states (Supplementary Fig. 13). Altogether, scTour allows simulation of cellular states that have not been captured during a scRNA-seq experiment.

### scTour can perform cross-platform, -system, -species predictions

Given the capability of scTour to characterise unseen cellular states, I next tested in a broader context the ability of scTour to predict the cellular dynamics of datasets that differ in many aspects from the one used to train the model. Here I selected the process of cortical excitatory neuron differentiation which has been well described in different species and biological systems using single-cell genomics^24–27^. Specifically, I trained the scTour model using a scRNA-seq dataset profiling the developing human cortex with the 3′ Kit v3 of 10x Genomics^24^. I analysed the same set of cells used in the original study for reconstruction of the excitatory neuron trajectory (36,318 cells). Before the model training, the excitatory neurons were relabelled according to their degree of maturity along the differentiation course (Supplementary Fig. 14a). The resulting scTour model, as expected, charted the cell differentiation trajectory from cycling progenitors, nIPCs, migrating neurons, immature to mature excitatory neurons, as evidenced by the developmental pseudotime, transcriptomic vector field and latent space robustly inferred, regardless of the substantial batch effects present in this dataset (Fig. 4a, Supplementary Fig. 14b).

**Fig. 4.**
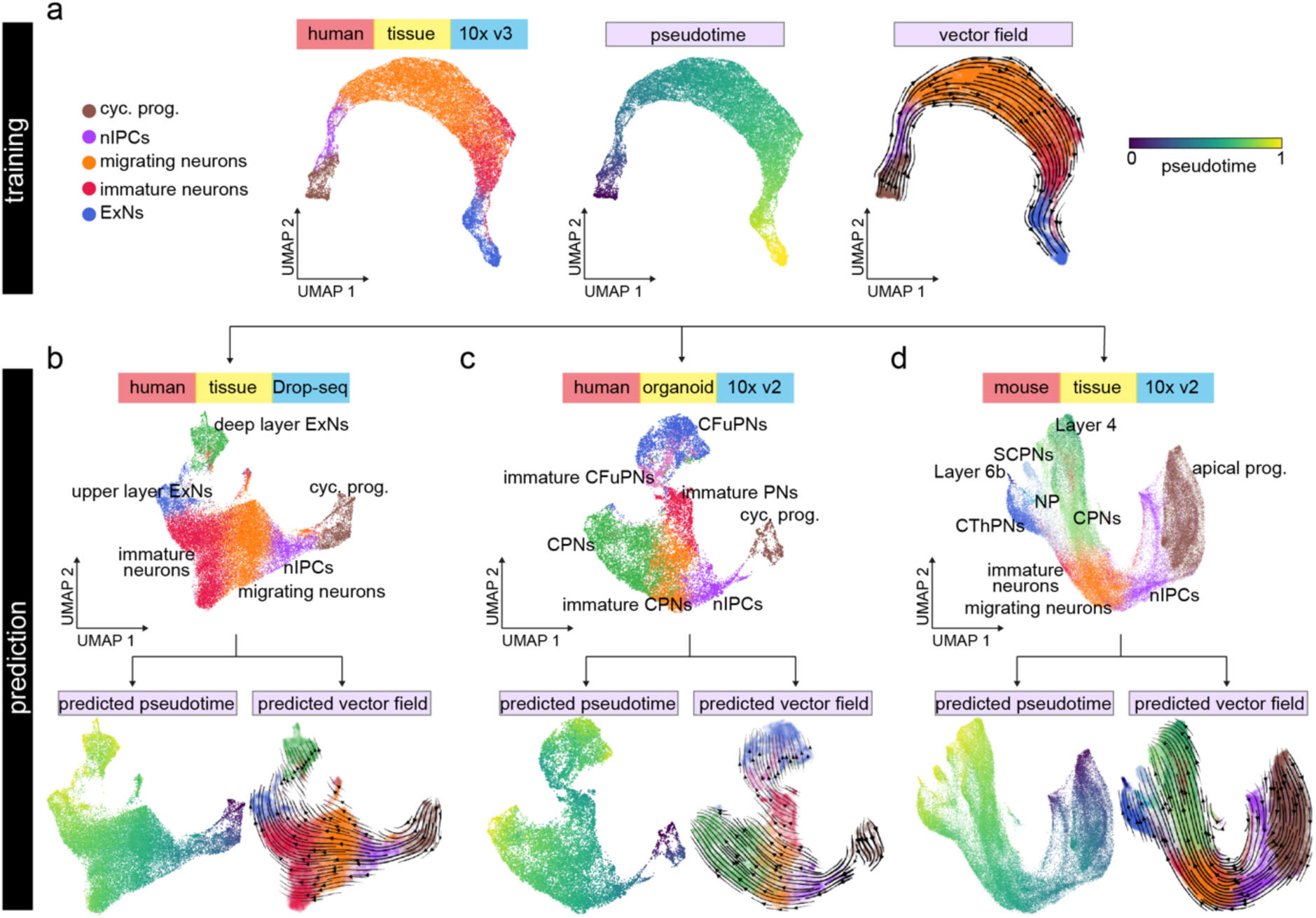
Cross-platform, -system, -species predictions of cellular dynamics during excitatory neuron development by scTour. **a,** UMAP visualizations of the latent space (left, coloured by cell types), developmental pseudotime (middle), and transcriptomic vector field (right) estimated by the scTour model trained using 60% of the 36,318 cells from the developing human cortex (10x Genomics)^24^. **b-d,** Upper: UMAP visualizations of the cell types from another developing human cortex dataset (Drop-seq, 27,855 cells)^25^ (**b**), a human brain organoid dataset (10x Genomics, 16,032 cells)^26^ (**c**), and a developing mouse cortex dataset (10x Genomics, 73,649 cells)^27^ (**d**). Bottom: the predicted pseudotime (left panels) as well as transcriptomic vector fields (right panels) for these three test datasets by the scTour model from **a**. cyc. prog., cycling progenitors; nIPCs, neuronal intermediate progenitor cells; ExNs, excitatory neurons; PNs, projection neurons; CPNs, callosal projection neurons; CFuPNs, corticofugal projection neurons; CThPNs, corticothalamic projection neurons; NP, near projecting; SCPNs, subcerebral projection neurons; apical prog., apical progenitors.

Given this model, I next assessed its performance in cross-data predictions by testing three additional datasets covering different experimental platforms, biological systems, and species: (1) Drop-seq-based measuring of the developing human cortex^25^ (27,855 cells); (2) an *in vitro* organoid system modelling the human cerebral cortex^26^ (10x Genomics 3′ Kit v2, 16,032 cells); (3) developing cortex from a different species, mouse^27^ (10x Genomics 3′ Kit v2, 73,649 cells). Despite large discrepancies between these three test datasets and the one used for training, scTour successfully reconstructed the cell trajectories mirroring excitatory neuron differentiation for all three datasets. This was shown by the precisely predicted pseudotime, vector field, and latent space without any prior corrections of batch effects present across all datasets (Fig. 4b-d and Supplementary Fig. 14c-e). Altogether, the dynamic properties of a new dataset can be efficiently decoded by scTour with a negligible time cost in prediction. It is thus a new useful tool for cross-data integrations and comparisons.

### Comparison of scTour with existing algorithms

A clear feature distinguishing scTour from currently available algorithms is its ability to jointly infer the pseudotime, vector field and latent representations of cells, as well as to predict cellular dynamics of unobserved data (Fig. 5a). To benchmark scTour against widely used methods, I assessed each of these functionalities separately (excluding the prediction functionality which is not available in other tools). Specifically, scTour was compared with scVelo^8^, Palantir^28^, Monocle 3^29^ and Slingshot^30^ for pseudotime estimation, with scVelo’s stochastic and dynamical models for vector field delineation, and with scVI^31^ for latent space inference. The benchmarking was conducted based on the process of excitatory neuron development as illustrated above^24^. This process has well-described ground truth for pseudotime and vector field comparisons and the data displayed significant batch effects for latent space assessment (36,318 cells, Supplementary Fig. 14b).

**Fig. 5.**
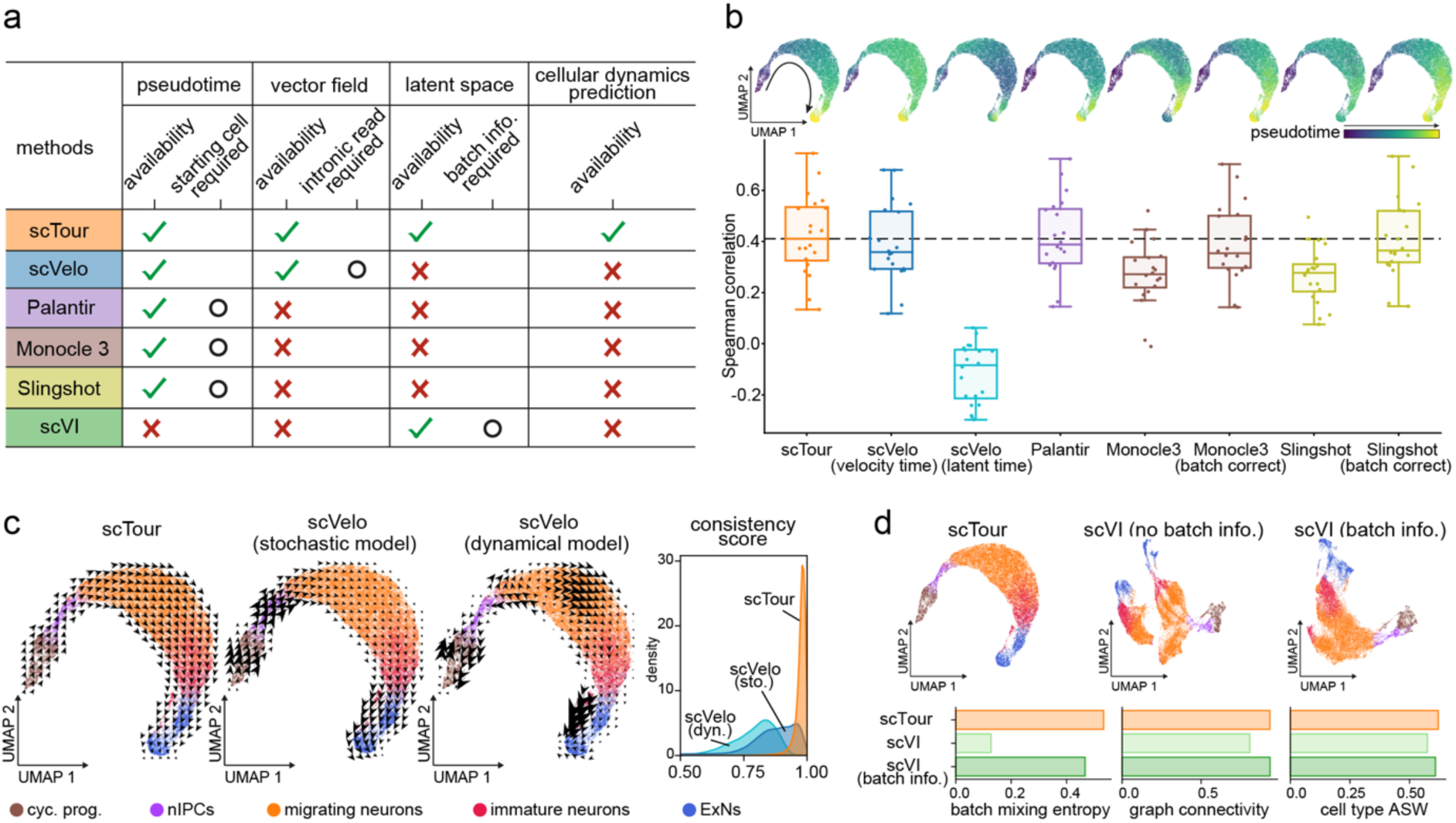
Benchmarking scTour against existing methods. **a,** Table summarizing the functionalities and features of scTour versus other methods. **b,** Upper panels: UMAP visualizations of pseudotime estimated by different algorithms along the excitatory neuron development (36,318 cells^24^). Lower panel: Box plot showing the Spearman correlation coefficients calculated between the pseudotime estimates and the expression profiles of 20 well-established marker genes along the excitatory neuron developmental trajectory (see Methods). The horizontal dotted line denotes the median value from scTour. **c,** UMAP visualizations of the vector fields from scTour (left), scVelo’s stochastic (middle) and dynamical (right) models, with colours indicating different cell types. The rightmost panel illustrates the distributions of the consistency scores calculated between each cell and its neighbouring cells based on the vector fields derived from each method. **d,** Upper panels: UMAP visualizations based on the latent space from scTour (left), scVI without batch correction (middle), and scVI with batch information incorporated during model training (right). Cells are colour coded by cell types as in **c**. Lower panels: metrics quantifying the degree of batch correction (batch mixing entropy and graph connectivity) and biological signal conservation (cell type ASW) for each method. ASW, average silhouette width; cyc. prog., cycling progenitors; nIPCs, neuronal intermediate progenitor cells; ExNs, excitatory neurons.

Comparison of the pseudotime estimated by different tools highlighted the superiority of scTour in several aspects. Firstly, scTour more accurately recapitulated the continuous progression from cycling progenitors to mature excitatory neurons than did other methods, as evidenced by the higher correlation between the pseudotime estimates and the expression patterns of well-established marker genes along the trajectory (Fig. 5b). Secondly, scTour has no demand for the starting cells which is required by Palantir, Monocle 3 and Slingshot. Although scVelo has no such requirement, the resulting pseudotime including the velocity pseudotime and latent time were not as accurate as that from scTour (Fig. 5b). Thirdly, the batch effects inherent in this data had no influence on scTour, but greatly impacted Monocle 3 and Slingshot as their performance dropped dramatically when batch correction was not performed prior (Fig. 5b).

At the level of the transcriptomic vector field, scTour was shown to both capture the underlying cellular dynamics and display considerably high consistency across neighbouring cells (Fig. 5c). By contrast, the stochastic model from scVelo exhibited much lower consistency scores and its dynamical model erroneously directed migrating neurons towards progenitor cells (Fig. 5c).

With respect to the latent space, scTour was compared with scVI which likewise yielded latent representations of cells. As expected, without providing the batch information during model training, scTour was able to largely alleviate the influence of these batches and meanwhile preserve the intrinsic biological signals, as illustrated by the latent space-based UMAP visualisation and by assessment of the batch mixing and biological conservation (Fig. 5d). By contrast, when the batch information was not considered and incorporated into the scVI model, the resulting latent space was dominated by sample batches, with cells from the same cell type segregated greatly across batches (Fig, 5d). Only after the batch factor was taken into account during modelling can scVI achieve the performance comparable to scTour (Fig. 5d).

### Characterization of human skeletal muscle development using scTour

To further showcase the biological insights that can be delivered through using scTour, I applied scTour to a challenging time series dataset which profiled the human limb muscle tissues over development from embryonic to adult stages (embryonic (prenatal weeks 5-8), fetal (prenatal weeks 9-18), juvenile (postnatal years 7-11) and adult (postnatal years 34-42))^32^. In the original study, cells from each stage were analysed separately, impeding the delineation of the whole developmental picture. Given scTour’s insensitivity to batch effects, it is possible to achieve the unbiased integration of cells across different time points to chart a biology-driven developmental trajectory. As a result, scTour reconstructed the full picture of human skeletal muscle ontogeny, with the diverse cell types chronologically placed in the low dimensional space and the pseudotime estimates in keeping with the real developmental time (Fig. 6a-c). This trajectory also showed that the non-myogenic cell populations including mesenchymal, chondrogenic and dermal fibroblast cells segregated between embryonic and fetal stages, indicating a pronounced transcriptomic change at this time window (Fig. 6a,c). The skeletal muscle (SkM) cells, on the other hand, displayed continuous changes during prenatal development, with major transcriptomic changes occurring postnatally (Fig. 6a,c).

**Fig. 6.**
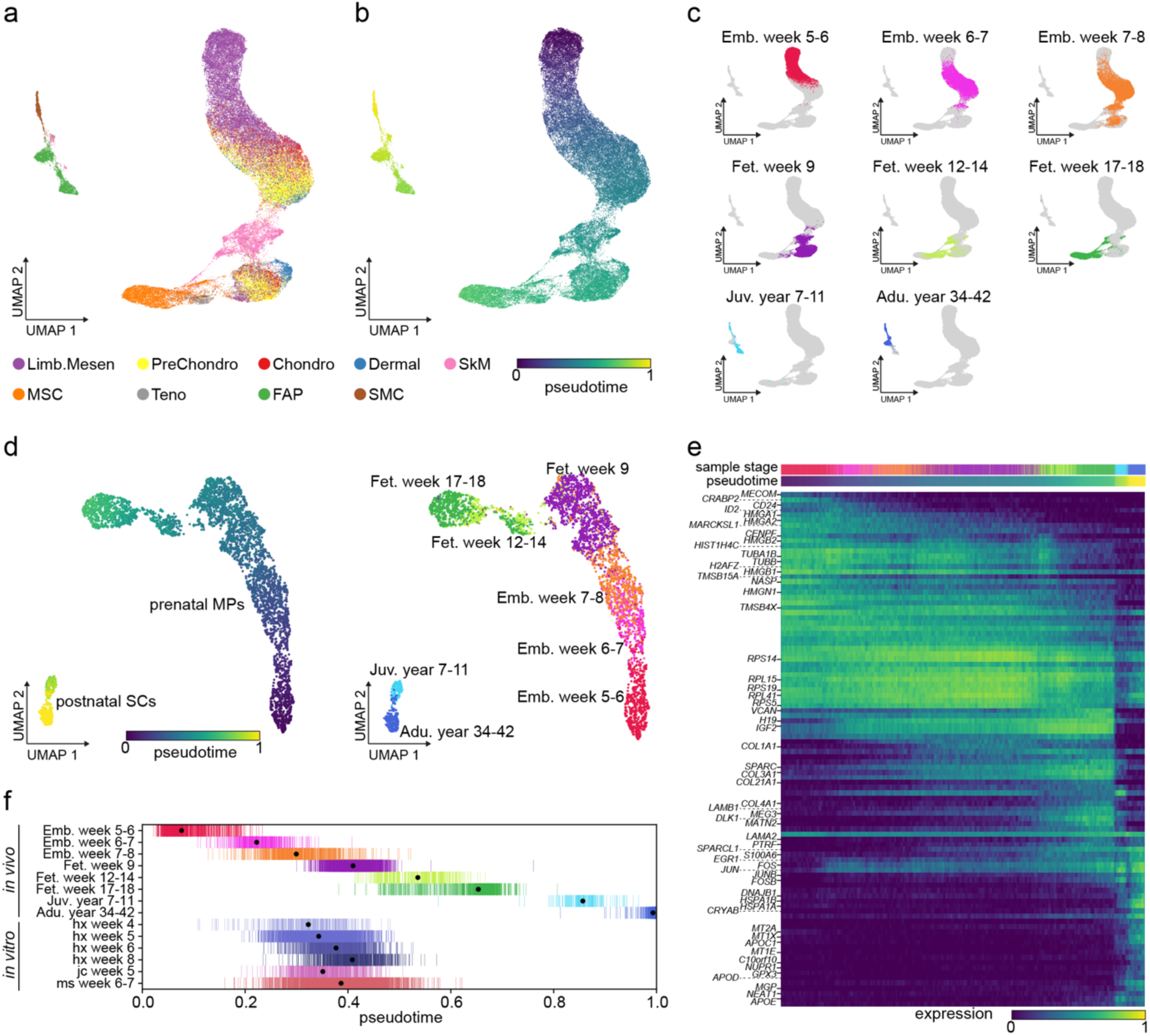
Application of scTour to human skeletal muscle development. **a,** scTour’s latent space-based UMAP visualization of cell types from all stages during human limb development (58,021 cells)^32^. **b,** As in **a**, but coloured by the pseudotime estimated by scTour. **c,** UMAP plots displaying cells from each of the developmental stages represented by different colours. **d,** scTour’s latent space-based UMAP representations of the skeletal muscle progenitor and stem cells collected across prenatal and postnatal development (4,816 cells), with colours indicating pseudotime estimates from scTour (left) and real developmental stages (right). **e,** Heatmap illustrating the expression dynamics of top 100 most significant genes along the developmental trajectory. Developmental stages and estimated pseudotime are displayed on top. **f,** Ordering of the *in vivo* skeletal muscle progenitor and stem cells based on the inferred pseudotime (upper), and ordering of the *in vitro* hPSC-derived progenitors based on the predicted pseudotime (lower). Dots denote the median values. Colours represent the developmental stages as in **d** (upper) and directed differentiation protocols and time points (lower). Limb.Mesen, limb mesenchymal cells; PreChondro, prechondrogenic cells; Chondro, chondrogenic cells; Dermal, dermal fibroblasts; SkM, skeletal muscle cells; MSC, mesenchymal stromal cells; Teno, tenogenic cells; FAPs, fibro-adipogenic progenitors; SMC, smooth muscle cells.

One of the major directions the original study sought to explore was the molecular changes of the skeletal muscle progenitor and stem cells along development. By using scTour to particularly analyse those cells including the prenatal myogenic progenitors (MPs) and postnatal satellite cells (SCs), the gradual transcriptomic changes for MPs during prenatal development and the evident separation between MPs and SCs were revealed (Fig. 6d). There was a gap in the trajectory between cells from fetal week 9 and those from weeks 12-14, possibly corresponding to the missing sample stage in between (Fig. 6d). Different from the previous work where the progenitor and stem cells were divided and assigned to five developmental stages, the unbiased pseudotime estimates from scTour allowed the investigation of the continuous molecular changes underlying the developmental progression.

Indeed, regression analysis of the gene expression changes along pseudotime identified previously uncovered transcriptional patterns underpinning the cellular dynamics (Fig. 6e). For instance, genes related to cell differentiation including *MECOM*, *CRABP2*, *CD24*, and *ID2*^33–36^ were specifically present in the earliest MPs (embryonic weeks 5-7) (Fig. 6e). Moreover, the three long noncoding RNAs identified (*H19*, *MEG3*, and *NEAT1*) were all involved in muscle differentiation^37^ and displayed distinctive dynamics: *H19* was mainly expressed from late embryonic to fetal stages while *MEG3* was more enriched in late fetal and juvenile stages, whereas *NEAT1* was exclusive to postnatal SCs (Fig. 6e). For the postnatal SCs, they showed high expression of immediate early genes (*EGR1*, *FOS*, *JUN*, *JUNB*, and *FOSB*) and genes encoding heat shock proteins (*HSPA1A*, *HSPA1B*, and *DNAJB1*), indicating the early activation of those SCs induced by the cell isolation procedure during the experimental collections^38^ (Fig. 6e). Besides, the SCs were enriched for genes associated with SC differentiation, regeneration and survival (*SPARCL1*, *CRYAB*, and *GPX3*)^39–41^, as well as genes involved in cell-cycle inhibition (*NUPR1*, *C10orf10/DEPP*)^42, 43^ to maintain their quiescence (Fig. 6e).

Another major question the original study aimed to answer was the developmental status of the *in vitro* skeletal muscle progenitor cells (SMPCs) that were derived from human pluripotent stem cells (hPSCs) through different directed differentiation protocols (HX, JC, MS)^32^. This was partially solved by aligning the *in vitro* SMPCs with the *in vivo* progenitor and stem cells in the diffusion map space, as well as by scoring each cell based on genes enriched in postnatal versus embryonic stages^32^. Such an analysis, however, mapped *in vitro* SMPCs to a broad time window, that was embryonic week 7 to fetal week 12^32^. Moreover, how the status of these cells varied across different protocols and directed differentiation time points were not clear. To address these questions, I used scTour to predict the developmental pseudotime of each *in vitro* SMPC on the basis of the model trained with *in vivo* SMPCs. This provided a higher-resolution view of the *in vitro*-*in vivo* alignment, and revealed the discrepancy among directed differentiation protocols used and time points collected (Fig. 6f). Specifically, by first focusing on the cells from 4-8 weeks of *in vitro* differentiation under the HX protocol, a clear trend arose: the directed differentiation process followed the patterns of *in vivo* development, with cells from shorter differentiation time matching an earlier *in vivo* developmental phase (Fig. 6f). For instance, cells undergoing 4 weeks of *in vitro* differentiation corresponded to the stage of embryonic weeks 7-8 while those with *in vitro* differentiation time of 8 weeks aligned to fetal week 9 (Fig. 6f), providing a finer time window (7-9 weeks) compared to the original study (7-12 weeks). Further comparison of the *in vitro* SMPCs obtained from different protocols showed a high similarity between the JC and HX protocols, with cells derived from them (both collected at week 5) consistently mapping to the *in vivo* stage of 7-9 weeks (Fig. 6f). Different from them, the MS protocol yielded more heterogeneous populations at 6-7 weeks of *in vitro* differentiation, spanning a broader developmental period from embryonic weeks 7-8 to fetal weeks 12-14 (Fig. 6f).

## Discussion

scTour is an innovative and comprehensive method for dissecting cellular dynamics by analysing datasets derived from single-cell genomics. It provides a unifying framework to depict the full picture of developmental processes from multiple angles including developmental pseudotime, vector field and latent space, and further generalises these functionalities to a multi-task architecture for within-dataset inference and cross-dataset prediction of cellular dynamics in a batch-insensitive manner.

There are several unique features of scTour compared to existing methods. In general, unlike the current algorithms which rely on either a batch-corrected low-dimensional space to estimate pseudotime, or an existing batch-corrected embedding to visualize the velocity field, scTour starts from the raw gene expression matrix and ends with the full developmental dynamics revealed under a single framework. The resulting latent space, which is not available in many trajectory inference tools, offers information on trajectory reconstructions, cell stratifications and data integrations. More importantly, all the inferences from scTour are invariant to batch effects, and the ultimate estimates are dominated by intrinsic biological signals. This presents a fascinating feature for exploring the cellular dynamics by integrating datasets from different studies, experimental platforms and systems. scRNA-seq data integration has been a challenging task and scTour provides an easy way to achieve this goal under the context of analysis of various dynamic processes.

scTour also introduces an alternative way to calculate transcriptomic vector fields. Compared to the state-of-the-art RNA velocity^7, 8^, scTour delivers several superiorities: (1) scTour does not require quantification of spliced and unspliced mRNAs, a rate-limiting but essential step in estimation of RNA velocity. (2) in scTour, delineating the developmental processes using pseudotime and transcriptomic vector field is highly convergent, as both are derived from the same model. In contrast, in scVelo, the RNA velocity field and the latent time are sometimes unmatched, presumably due to the different strategies taken when summarising gene-wise estimates into cell-level inferences^8^. (3) RNA velocity estimates can be affected by genes with partial or no kinetics captured^9, 11^. This has no impact on scTour’s vector field (Supplementary Fig. 9). (4) the application of RNA velocity to single-cell epigenetic data is not feasible and to single-nucleus data is limited, due to the need to model transcriptional kinetics using spliced and unspliced reads. scTour overcomes these limitations as it relies only on the abundance matrix which quantifies the amount of transcripts/chromatin accessibility across cells. It is thus applicable to datasets of both snRNA-seq (Supplementary Fig. 8) and scATAC-seq (Supplementary Fig. 15). (5) scTour’s vector field can be predicted based on the learned differential equation for a new dataset agnostic to the scTour model, a feature not available in scVelo. All these features broaden the use of vector field to decode dynamic processes with scTour.

The uniqueness of scTour also lies in its prediction functionalities comprising predicting cell characteristics given the transcriptomes and predicting the transcriptomic latent space given the time interval. This prediction is robust across biological systems, species and experimental platforms, and provides a convenient way for cross-data comparisons by propagating the information from existing datasets to new ones.

In this study scTour’s new features and usefulness are obvious in multiple datasets. Given its robust performance with respect to batch effects and ability to scale to large datasets, I anticipate scTour will be of immediate interest to a broad community of users.

## Methods

### The scTour model

scTour models the cellular dynamics under the framework of VAE^13^ and neural ODE^14^. By taking as input an abundance matrix (e.g. a gene expression matrix with *n* cells and *g* genes) *x* ∈ *R^n×g^*, a probabilistic encoder network *f_z_* is used to approximate the posterior *q*(*z*|*x*) by assuming a multivariate Gaussian with a diagonal covariance, with the mean *μ* and standard deviation *σ* of the approximate posterior generated from *f_z_*. *z* is then sampled from *q*(*z*|*x*) through the reparameterization trick^13^:

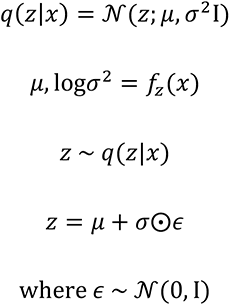

A second encoder network *f_t_*, with the hidden layer shared with *f_z_*, transforms *x* into a scalar time *t* in the 0-1 range through the Sigmoid function. This corresponds to the developmental pseudotime of a given cell. By sorting cells based on their time *t*, the latent state *z* at *t*_0_ can be obtained. Next, given the initial state *z_t_*_0_ and times *t*_0_, *t*_1_, *t*_2_, …, *t_n_* across cells, an ODE solver generates *z_t_*_1_, *z_t_*_2_,…, *z_t_*_n_ based on the differential equation (the derivative of the latent states with respect to time) which is defined by another neural network *f_ode_*:

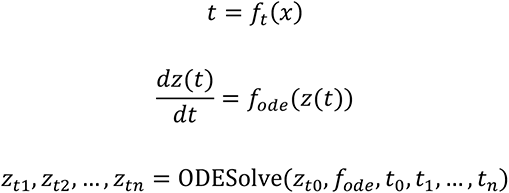

The “odeint” function within torchdiffeq^14^ is used to perform this task.

Subsequently, the latent *z* sampled from the approximate posterior, and the *z_t_* from the ODE solver parallelly go through a decoder network *f_d_* to reconstruct *x*. The objective function here is a modified lower bound:

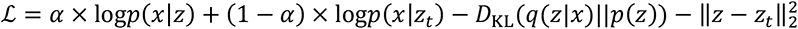

where the prior *p*(*z*) is the standard multivariate Gaussian here. This equation combines the weighted reconstruction errors from both *z* and *z_t_*, the Kullback–Leibler divergence of the approximate posterior from the prior, and the mean squared error (MSE) between *z* and *z_t_* as a regularizer to tune *z_t_* towards *z*.

In scTour, there are three modes to calculate the reconstruction errors, namely, MSE, negative binomial (NB)-conditioned likelihood and zero-inflated negative binomial (ZINB)- conditioned likelihood. MSE is a straightforward metric to measure the distance between the reconstructed and observed *x*. This mode requires the log-transformed normalized expression matrix as the input, and exhibits good performance with much less runtime cost. Compared to the MSE mode, NB mode relies on the raw count matrix and assumes a NB distribution for *p*(*x*|*z*), with the gene dispersion parameter estimated by an additional neural network. Besides, similar with the scVI model^31^, the decoder network outputs the abundance proportion of each gene in a given cell via the softmax activation. The final reconstructed expression is obtained through multiplying this proportion by the library size which is approximated by summing the raw counts across genes within a cell here. ZINB mode requires the raw count matrix as in the NB mode, but models the gene expression based on the assumption of a ZINB distribution. Additionally, it uses a neural network to compute the dropout probability as in scVI.

All the hidden layers use ReLU as the activation function except for the neural net *f_ode_* where ELU is used.

### Model training, inference, and prediction

#### Model training

Although the application of mini-batches in neural ODE is less straightforward^14^, mini-batch training suits the scTour architecture quite well, which offers a number of advantages. Specifically, mini-batch training makes direct backpropagation more feasible, model training faster, and memory more efficient. These together endow scTour with the great scalability to large datasets. Importantly, with mini-batch training, scTour is able to achieve high performance using only a subset of cells sampled. The batch size is set to 1,024 throughout the paper and can be adjusted depending on datasets. For the optimization, scTour uses Adam as the optimizer, with the L2 regularization implemented to strengthen model generalisation. Since scTour converges faster for large datasets versus small ones, the default number of epochs in scTour is proportional to the number of cells in the dataset of interest.

#### Subsampling-based training

scTour provides the option to train the model with a subset of cells. Specifically, scTour first shuffles the entire dataset and then randomly samples a given proportion of cells from the shuffled data. The two rounds of randomness ensure the preservation of the cellular diversity. This step reduces the training time and has marginal influence on the model performance as shown in multiple datasets.

#### Cellular dynamics inference

After the model training, scTour assigns a developmental pseudotime to each cell based on the learnt time neural net *f_t_* without the need for specifying starting cells. Since there exist two possible integration directions (forward or backward), the inferred pseudotime can be in the correct ordering (ascending), or the reverse (descending). To resolve this, scTour leverages the information of gene counts (i.e., the number of expressed genes) across cells which is demonstrated to anti-correlate with developmental potential^44^. Specifically, a linear regression model is fit between the inferred pseudotime and the gene counts. If the slope is positive, the estimated time will be reversed, and the downstream predictions will be reversed as well. In the cases where the use of gene counts fails to capture the expected trend, scTour provides a post-inference function to reverse the pseudotime.

The transcriptomic vector field is the learnt differential equation *f_ode_*, which outputs the gradient given the current latent state and thus provides information regarding the future transcriptomic directions.

The latent representations of cells in scTour are the weighted combination of *z* from the variational inference and *z_t_* from the ODE solver:

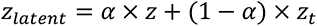

Larger *α* skews the latent space towards the intrinsic transcriptomic structure while smaller *α* is more representative of the extrinsic pseudotime ordering. Users of scTour have the option to adjust *α* according to their purposes.

#### Cellular dynamic prediction

Given the gene expression matrix of query cells from an unobserved cellular state or a new dataset, scTour predicts their developmental pseudotime by the time neural net *f_t_*, transcriptomic vector field by the function *f_ode_*, and latent representations by the whole framework built from reference cells.

Regarding the prediction of the transcriptomic space given an unobserved time interval *t*_1,_ *t*_2_,…, *t_n_*, scTour takes a stepwise integration given the learnt differential equation *f_ode_* by leveraging the *k*-nearest neighbours. Specifically, the developmental pseudotime *T* and the latent representations *Z* from the training data are used as a reference. Next, for each time point *t* within the unobserved interval, its *k*-nearest neighbours in the reference are obtained by comparing *t* with *T*. Next for each neighbour *j*, the ODE solver takes the latent state of this neighbour *z_j_* as the initial value, together with the time of this neighbour *t_j_* and the time *t*, to output the latent state corresponding to *t*. The final latent representation of the time *t* is calculated as the average across the *k*-nearest neighbours:

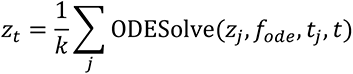

For each time point estimated, the resulting latent state *z_t_* along with the time *t* are added to the latent state *Z* and time *T* pool to update the reference for predicting the next time point. This procedure is stopped until the entire time span has been predicted.

### Visualization of vector field

The visualization of the transcriptomic vector field on a low-dimensional embedding such as UMAP is obtained using a similar approach as in velocyto^7^ and scVelo^8^. The main idea is to position the velocity arrow in the direction where the estimated velocity best matches the transcriptomic difference. To this end, a cell-cell transition probability matrix *P* is first calculated. Different from velocyto and scVelo which calculate this matrix using the gene-based velocity vector and the gene expression difference, scTour computes the matrix at the level of latent space. Specifically, based on the vector field derived from the learnt differential equation *f_ode_* and the latent state of each cell, scTour calculates the cosine similarity between the gradient and the latent difference:

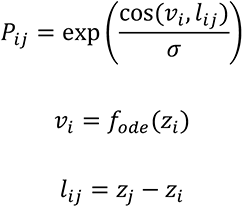

where *ν*_1_ is the gradient of cell *i* inferred from the learnt differential equation *f_ode_* given its latent state *z_i_*, and *l_ij_* represents the difference between cell *i* and *j* at the latent space level. Both *ν*_1_ and *l_ij_* can be optionally transformed using variance-stabilizing transformations before calculating the cosine similarity. Similar with scVelo, for each cell, only the recursive neighbours from the KNN graph are considered for cell-cell transition probability estimation. Differently, scTour also considers the neighbours in the time space based on the developmental pseudotime inferred for each cell. The resulting transition probability matrix *P* is next row-normalized to let ∑*_j_P_ij_* = 1. The normalized matrix is used as weights to calculate the unitary displacement vector for each cell:

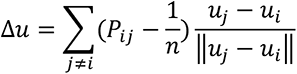

where *μ_i_* and *μ_j_* are the coordinates of cells *i* and *j* in the low-dimensional embedding. This displacement vector can be visualized for each cell or on the grid level as arrows or streamlines.

### Analysis of mouse dentate gyrus neurogenesis

The two datasets from mouse dentate gyrus used in Fig. 2 are from^15^. For the first dataset, the raw count matrix and meta information were downloaded from Gene Expression Omnibus (GEO) under the accession number GSE95315. Only the cell types along the granule cell lineage including nIPCs, neuroblasts (Neuroblast_1, Neuroblast_2), immature and mature granule cells were used for the following analysis (4,007 cells). Before running scTour, the data was preprocessed by filtering genes detected in less than 20 cells and selecting the top 500 highly variable genes using Scanpy^45^. A scTour model was then trained with the raw count matrix from these 500 genes across 4,007 cells. The resulting model was used to infer the developmental pseudotime, transcriptomic vector field and latent representations of these cells (the latent space was generated with 20% *z* and 80% *z_t_*). UMAP embeddings derived from the inferred latent space and PCA space (40 PCs) were compared. For the comparison of the vector field between scTour and scVelo in Supplementary Fig. 1, the cells from the two time points P12 and P35 which were used in the scVelo publication were extracted to run scTour and scVelo.

For the second dataset downloaded from GEO (GSE104323), the cells from the granule lineage (nIPCs, neuroblast, immature and mature granule cells) and the pyramidal lineage (immature pyramidal cells) were considered (15,174 cells). Similarly, genes detected in less than 20 cells were excluded and the top 2,000 highly variable genes were used for scTour model training, which yielded the developmental pseudotime, vector field and latent space (40% *z* and 60% *z_t_*) of cells. The latent space from scTour and PCA space (30 PCs) were used to calculate the UMAP embeddings for comparisons. To demonstrate the robustness of scTour model to cell subsampling, the models were trained based on cell subsets from 1% to 95% of all cells. The resulting models were used to infer the dynamics (developmental pseudotime, vector field, and latent representations) of all cells. Spearman correlation coefficients between the developmental pseudotime derived from the models trained with <95% of all cells and that from the model trained with 95% of cells were calculated to show the stable inference.

### Analysis of mouse pancreatic endocrinogenesis

The dataset from mouse pancreatic endocrine development^8, 23^ used in Fig. 3 was downloaded from scVelo package. The scTour model training started from the raw count matrix including the top 2,000 highly variable genes and 3,696 cells, and ended with the estimated developmental pseudotime, transcriptomic vector field and latent representations (70% *z* and 30% *t_z_*) of the cells. To obtain the latent time from scVelo’s dynamical model, the same procedure as in the original scVelo publication was used to reproduce the results.

To test the ability of scTour to predict the dynamics of unseen cellular states, the model was trained by excluding one of the cell types and the resulting model from the remaining cell types was used for two purposes: (1) predicting the developmental pseudotime, transcriptomic vector field, and latent representation of the excluded cell type given its gene expression matrix; (2) predicting the latent representation of the excluded cell type given its expected developmental time along the differentiation path. The comparison of the predicted latent representation with the ground truth (the latent space of the excluded cell type derived from its gene expression matrix) was performed from three angles. Firstly, the predicted latent space, together with the latent space of all cell types during endocrinogenesis, were combined to yield a UMAP embedding. Secondly, the pairwise Euclidean distance was calculated between the predicted latent representation and the latent representation of each cell type. Lastly, unsupervised hierarchical clustering was conducted based on the predicted latent space and the latent space of all the cell types (Euclidean distance as the distance metric and ‘ward’ as the linkage algorithm).

### Analysis of cortical excitatory neuron development

Datasets profiling cortical excitatory neuron development used in Fig. 4 were from four sources: (1) the developing human cortex measured using 3′ Kit v3 protocol of 10x Genomics^24^. Here I focused on the same set of cells which were used in the original study to reconstruct the excitatory neuron developmental trajectory (36,318 cells). (2) the developing human cortex measured using Drop-seq^25^, with the cell types of cycling progenitors, intermediate progenitors, migrating neurons, maturing neurons, upper and deep layer excitatory neurons (27,855 cells) considered here. (3) the human brain organoid measured using 3′ Kit v2 of 10x Genomics^26^. Here I focused on the cells of cycling progenitors, intermediate progenitors, immature and mature excitatory neurons from the organoids cultured for three months (three-month PGP1 organoids 1-3, 16,032 cells). (4) the developing mouse cortex measured using 3′ Kit v2 of 10x Genomics^27^. The cells of apical progenitors, intermediate progenitors, migrating neurons, immature neurons, and excitatory neurons from different layers with different projection properties (73,649 cells) were used.

For the first dataset, since the excitatory neuron subtypes in the original study were labelled with arbitrary numbers, I relabelled those cells according to the second dataset where the excitatory neuronal cells were named on the basis of their maturity along the differentiation path. Specifically, CellTypist^46^ was used to train a model based on the reference dataset (i.e., the second one), which was subsequently employed to transfer the cell type labels to cells of the first one.

The scTour model was then trained based on the first dataset (training data) by using 60% of all cells, and 765 genes which were the intersection of the top 1,000 highly variable genes from this data with the genes detected in all the other three datasets (test data). This model was used to infer the developmental pseudotime, transcriptomic vector field, and latent space (50% *z* and 50% *z_t_*) of the training data (Fig. 4a), and to predict the properties of cells from the test data (Fig. 4b-d). For the UMAP embeddings of the three test datasets shown in Fig. 4b-d, the first two were derived from PCA-space (30 PCs) and the last one was batch corrected using BBKNN^47^ to mitigate the substantial batch effects among donors. For the UMAP embeddings of the three test datasets shown in Supplementary Fig. 14c-e, they were all derived from the predicted latent space by scTour without any batch corrections.

### Analysis of other biological processes

In addition to the developmental courses mentioned above, scTour was applied to a number of dynamic biological processes described as follows.

#### Mouse gastruloid

this dataset (30,496 cells) came from a study on embryonic gastruloid measured using 10x Genomics^16^. The cell type classification and UMAP embedding from the original study were used as is here. The developmental pseudotime, transcriptomic vector field, and latent representations (70% *z* and 30% *z_t_*) of these cells were inferred from the scTour model which was trained with 2,000 highly variable genes and 60% of cells randomly sampled from the whole data.

#### Human thymic epithelial cell development

this dataset (14,217 cells) profiled the human thymic epithelial development using 10x Genomics^17^. The cell annotations and UMAP embedding from the publication were used as is. The highly variable genes from the original study (804) and cells randomly sampled from the whole data (60%) were used to train the scTour model, which generated the developmental pseudotime, transcriptomic vector field, and latent representations (70% *z* and 30% *z_t_*) of all cells.

#### Human gastrulation

this dataset (1,195 cells) was from a gastrulating human embryo measured using Smart-seq2^18^. The cell annotations and UMAP embedding from the original study were used here. For scTour model training, the top 2,000 highly variable genes were considered. The trained model was then used to infer the developmental pseudotime, vector field, and latent representations (80% *z* and 20% *z_t_*) for these cells.

#### Human preimplantation

this dataset has 90 cells from human preimplantation embryos with single cells isolated by mouth pipette^19^. For the PCA-based UMAP embedding, the top 30 PCs derived from the 2,000 highly variable genes were used. The developmental pseudotime, transcriptomic vector field, and latent representations (70% *z* and 30% *z_t_*) of these cells were inferred from the scTour model trained with the same set of genes.

#### Reprogramming in mouse

this dataset (251,203 cells) was from a time course of iPSC reprogramming measured using 10x Genomics^20^. The original cell annotations and force- directed layout embedding (FLE) from the publication were used here. The scTour model was trained based on 2,000 highly variable genes and 20% of cells, which produced the developmental pseudotime, transcriptomic vector field, and latent representations (30% *z* and 70% *z_t_*) of all cells.

#### Reprogramming in human

this snRNA-seq dataset (36,597 nuclei) was from a study on human cell reprogramming^21^. Similarly, the cell annotations and UMAP embedding provided by the original study were used to visualize the estimated developmental pseudotime and transcriptomic vector field from the scTour model trained on the basis of 2,000 highly variable genes and 60% of all cells. The inferred latent space from the same model (70% *z* and 30% *z_t_*) was used to generate a new UMAP embedding to illustrate the reprogramming trajectory.

#### Human hematopoiesis

this scNT-seq dataset (1,947 cells) was from in vitro culture of the CD34+ human hematopoietic stem and progenitor cells (HSPCs)^9^. The gene set (1,956 genes) from the original study was used to train the scTour model, which yielded the pseudotime, transcriptomic vector field, and latent representations (80% *z* and 20% *z_t_*) of all cells. The cell annotations and UMAP embedding from the publication were used here for visualization.

#### Brain endothelial topography

this dataset (3,105 cells) was focused on the endothelial cells of the mouse brain^22^. To be consistent with the original study, the three subclusters (choroid plexus, artery shear stress, and interferon) were excluded from the differentiation trajectory reconstruction. The PCA space-based UMAP embedding was from the top 30 PCs which were obtained from the 2,000 highly variable genes. The trajectory reconstruction by scTour (developmental pseudotime, transcriptomic vector field, latent representations (20% *z* and 80% *z_t_*)) was based on the same set of genes.

#### Human fetal retinal chromatin accessibility

this scATAC-seq dataset (4,883 cells) was from human fetal retina^48^. Preprocessing of this dataset was conducted using Signac^49^, including normalization by term frequency-inverse document frequency (TF-IDF), feature selection (top 25% of peaks), and dimension reduction by singular value decomposition. The first latent semantic indexing (LSI) component which was highly correlated with sequencing depth was excluded and the 2-30 components were used for UMAP embedding calculation. The TF-IDF matrix (34,670 genomic regions across 4,883 cells) was used as the input for scTour model training, which generated the developmental pseudotime, epigenetic vector field, and latent representations (50% *z* and 50% *z_t_*) for these cells.

### Benchmarking scTour against existing algorithms

To benchmark scTour against existing methods including scVelo^8^, Palantir^28^, Monocle 3^29^, Slingshot^30^, and scVI^31^, the dataset profiling the developing human cortex by 10x Genomics (36,318 cells)^24^ was used. For all the analyses (pseudotime, vector field and latent space) performed by these tools, the top 1,000 highly variable genes were considered. The analytical procedure for each method is described as follows.

For scTour, 20% of cells randomly sampled from the entire dataset were used to train the scTour model, which yielded the developmental pseudotime, transcriptomic vector field, and latent representations (five dimensions, 50% *z* and 50% *z_t_*) of cells.

For scVelo, both the stochastic and dynamical models were performed following the tutorial at https://scvelo.readthedocs.io, with the velocity pseudotime derived from the stochastic model and the latent time derived from the dynamical model.

For Palantir, the pseudotime estimation was conducted based on the tutorial at https://nbviewer.org/github/dpeerlab/Palantir/blob/master/notebooks/Palantir_sample_notebook.ipynb. 40 PCs were considered during the diffusion map construction and the cell expressing the highest level of *PAX6* was designated as the starting cell when determining the pseudotime.

For Monocle 3, the Seurat Wrappers package was used to run Monocle 3 on the Seurat objects with or without batch effect correction, respectively. To obtain the batch-corrected Seurat object, the procedures from the tutorial at https://satijalab.org/seurat/articles/integration_introduction.html were followed^50^. The starting cell was specified as the one with the highest expression level of *PAX6* when estimating the pseudotime.

For Slingshot, the pseudotime was estimated based on a two-dimensional UMAP embedding (derived from 30 PCs) and a vector of clustering labels (from Louvain clustering at resolution of 0.2). The starting cluster was set to the one with the highest proportion of cycling progenitor cells. The final pseudotime was calculated as the average across the lineages. To rerun Slingshot by removing batch effects, the integration process from Seurat as described above was conducted to obtain batch corrected UMAP and clustering labels for pseudotime estimation.

For scVI, the scVI model training was performed using the default parameters to generate a 10-dimensional latent space. This was run twice, with or without batch information provided, respectively.

To examine the accuracy of the pseudotime estimated from different methods, the well-established marker genes along the excitatory neuron developmental trajectory were collected, including the markers for progenitors (*GLI3*, *TFAP2C*, *PAX6*, *SOX2*, *EOMES*, *JUND*, *NFE2L2*, *SOX9*, *EMX2*, and *FOS*) and excitatory neurons (*MEF2C*, *SATB2*, *STMN2*, *NEUROD2*, *NEUROD6*, *BHLHE22*, *POU2F2*, *ZBTB18*, *CHD3*, and *MYT1L*)^24, 25^. Spearman correlation coefficient was then calculated between the pseudotime estimates and the expression profiles of each of these genes.

To check the consistency of the vector field across neighbouring cells, consistency score, the same metric as defined in the scVelo publication, was computed here, which was the mean Pearson correlation coefficient calculated between the vector field of a given cell and its neighbours.

To evaluate the degree of batch correction and biological signal conservation, three metrics from scArches^51^ and scIB^52^ were used. The first one was the entropy of batch mixing, which measured the batch diversity in the neighbouring cells. 15 nearest neighbours were considered for each cell. The second metric was the graph connectivity, which estimated the connectivity among all the cells in each cell type. These two metrics were used to assess the batch correction. The third metric was the cell type average silhouette width, which calculated the inter-cluster (the nearest cluster) versus intra-cluster distances and was used to assess the biological signal conservation.

### Analysis of human skeletal muscle development

The scRNA-seq data profiling the human limb muscle tissues across prenatal and postnatal development, as well as the hPSC-derived *in vitro* muscle cells from different protocols was downloaded at skeletal-muscle.cells.ucsc.edu^32^. To investigate the full dynamics of human skeletal muscle ontogeny, a scTour model was trained using cells collected from all the developmental stages. The cell types with less than 1,000 cells (skin cells, red blood cells, Schwann cells, white blood cells, and endothelial cells) were excluded from the model training, resulting in 58,021cells as the input for scTour. The training was done based on the top 2,000 highly variable genes and 90% of all cells for 200 epochs, and the resulting model was used to infer the pseudotime and latent space (50% *z* and 50% *z_t_*) for all the cells.

Next, another scTour model was trained focusing on the skeletal muscle progenitor and stem cells (4,412 prenatal myogenic progenitors and 404 postnatal satellite cells). The training was conducted on the basis of 90% of all cells, and 1,791 genes which were the intersection of the top 2,000 highly variable genes with genes expressed in the *in vitro* skeletal muscle progenitors. The resulting model was used for two purposes: 1) to infer the pseudotime and latent space (50% *z* and 50% *z_t_*) of the *in vivo* progenitor and stem cells; 2) to predict the pseudotime of the *in vitro* progenitor cells derived from different directed differentiation time points and protocols (14,996 cells). To identify genes showing dynamic expression changes along the trajectory, the same method defined in Monocle^6^ was used. In detail, for each gene the cross-cell expression level was modelled as a function of the cells’ pseudotime by using cubic smoothing spline with three degrees of freedom in the R package VGAM^53^. The significance was estimated by the likelihood ratio test which compared the full model with the reduced model (i.e., intercept-only regression), with the *p*-value adjusted by the Bonferroni correction. The top 100 most significant genes were shown in Fig. 6e.

## Data availability

All the datasets used in this study are publicly available and summarized in Supplementary Table 1.

## Code availability

The source code of scTour is available at https://github.com/LiQian-XC/sctour.

## Acknowledgements

I thank A. Moffett for the support on this project and feedback on the manuscript.

## Author contributions

Q.L. conceived and implemented the scTour algorithm, and performed all the analyses and wrote the manuscript.

