## Supplementary Figs. 1-15 for "scTour: a deep learning architecture for robust inference and accurate prediction of cellular dynamics"

**Supplementary Information**

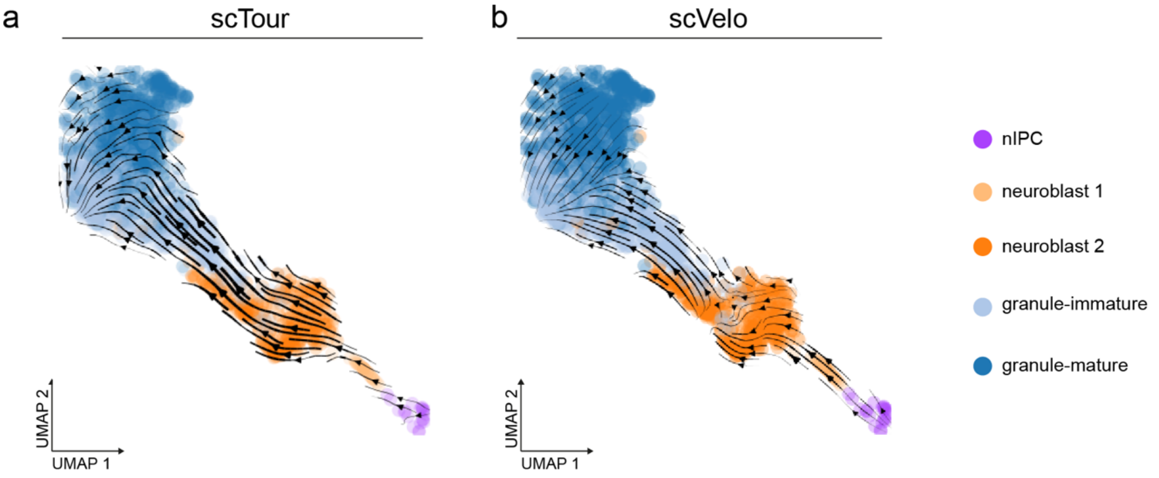


**Supplementary Figure 1 | Inferred vector field of cells during granule cell differentiation in dentate gyrus using scTour and scVelo.**

UMAP visualizations of the transcriptomic vector field inferred by scTour (**a**) and the RNA velocity inferred by scVelo (**b**) using the cells from the granule cell lineage. Both are inferred based on the cells from the two time points P12 and P35 as in the scVelo publication.

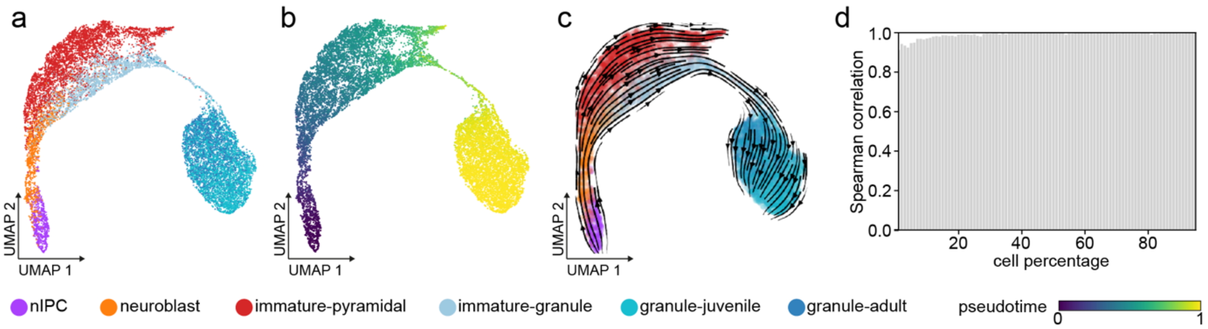


**Supplementary Figure 2 | scTour's inference is robust to cell subsampling.**

UMAP visualizations of the latent space (**a,** coloured by cell types), developmental pseudotime (**b**), and transcriptomic vector field (**c**) inferred from the scTour model trained using 1% of cells from the pyramidal and granule cell lineages. **d,** The Spearman correlation coefficient calculated between the pseudotime estimated from the model trained using 95% of total cells and those from models trained with cell subsets (1% to 95% from left to right).

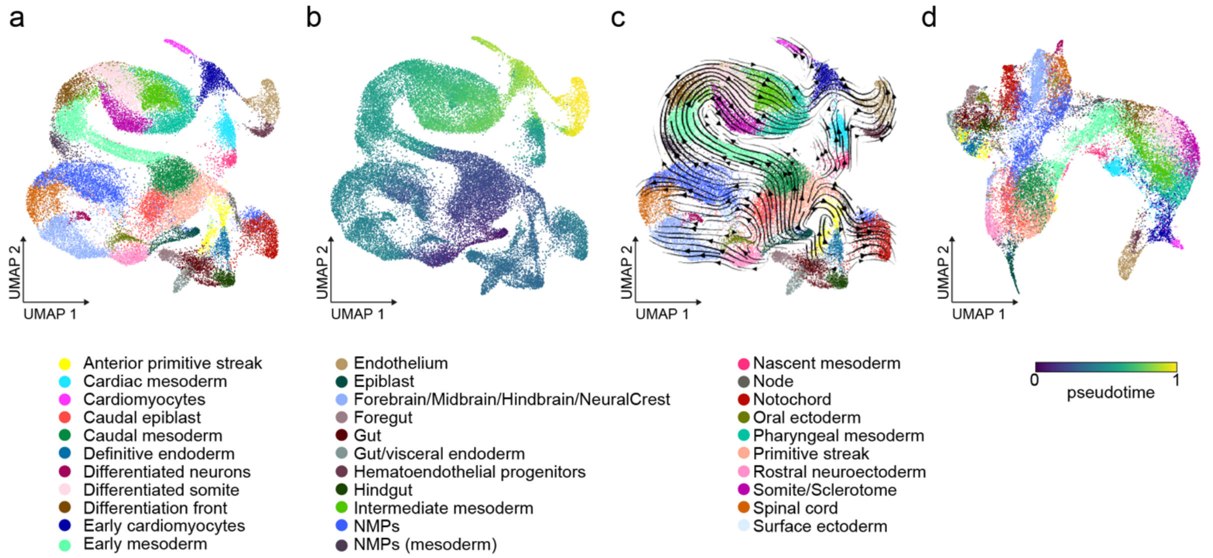


**Supplementary Figure 3 | scTour captures the developmental cellular dynamics in embryonic organoids.**

**a,** UMAP visualization of cell types from 30,496 cells collected from mouse gastruloids. The UMAP coordinates in **a-c** and cell type annotations are drawn from the original study. NMPs, neuromesodermal progenitors.

**b-d,** UMAP visualizations of the inferred developmental pseudotime (**b**), transcriptomic vector field (**c**), and latent space (**d**) from the scTour model trained using 60% of all cells.

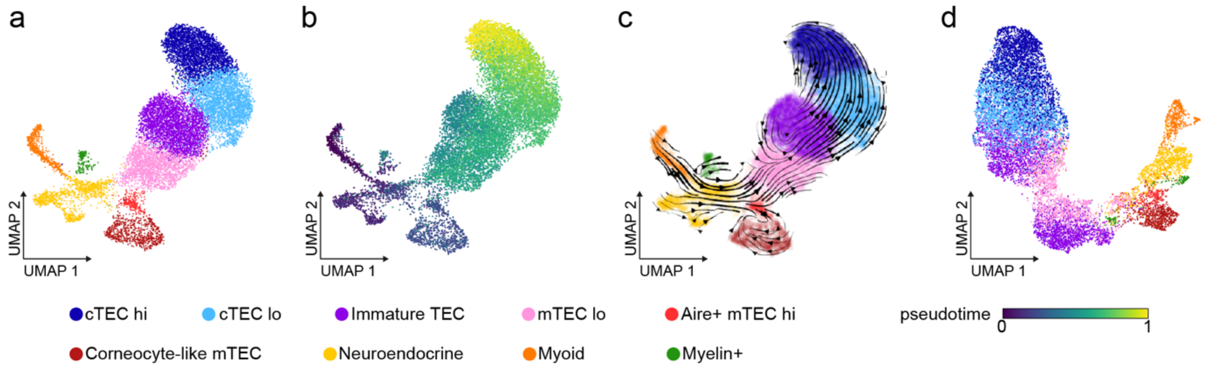


**Supplementary Figure 4 | scTour captures the developmental cellular dynamics in human thymic epithelial cells.**

**a,** UMAP visualization of cell types from 14,217 cells collected from human thymic epithelial compartment at different stages. The UMAP coordinates in **a-c** and cell type annotations are from the original study. TEC, thymic epithelial cells; cTEC, cortical TEC; mTEC, medullary TEC.

**b-d,** UMAP visualizations of the inferred developmental pseudotime (**b**), transcriptomic vector field (**c**), and latent space (**d**) from the scTour model trained using 60% of all cells.

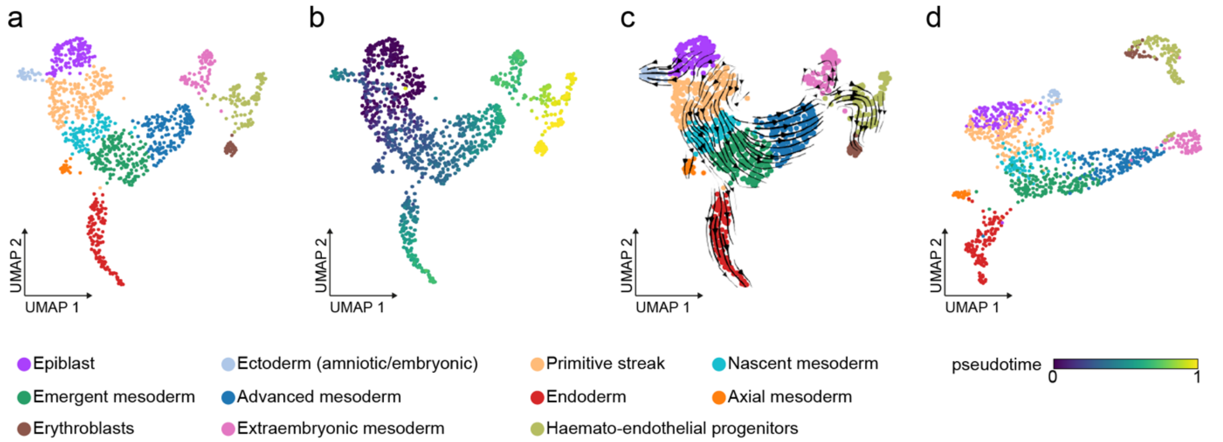


**Supplementary Figure 5 | scTour captures the developmental cellular dynamics in human gastrulation.**

**a,** UMAP visualization of cell types from 1,195 cells collected from a gastrulating human embryo. The UMAP coordinates in **a-c** and cell type annotations are from the original study.

**b-d,** UMAP visualizations of the inferred developmental pseudotime (**b**), transcriptomic vector field (**c**), and latent space (**d**) from the scTour model trained using 90% of all cells.

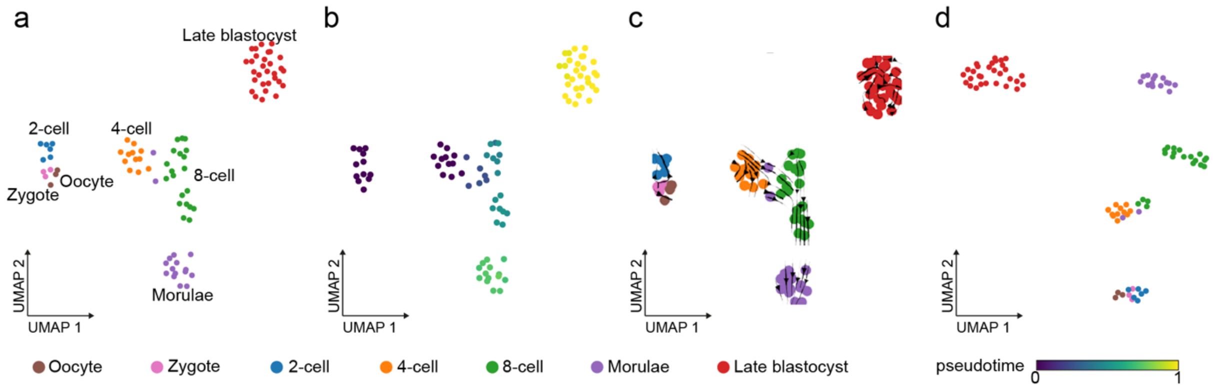


**Supplementary Figure 6 | scTour captures the developmental cellular dynamics in human preimplantation.**

**a,** PCA space-based UMAP visualization of the 90 cells collected from human preimplantation embryos.

**b-d,** UMAP visualizations of the inferred developmental pseudotime (**b**), transcriptomic vector field (**c**), and latent space (**d**) from the scTour model trained using 90% of all cells.

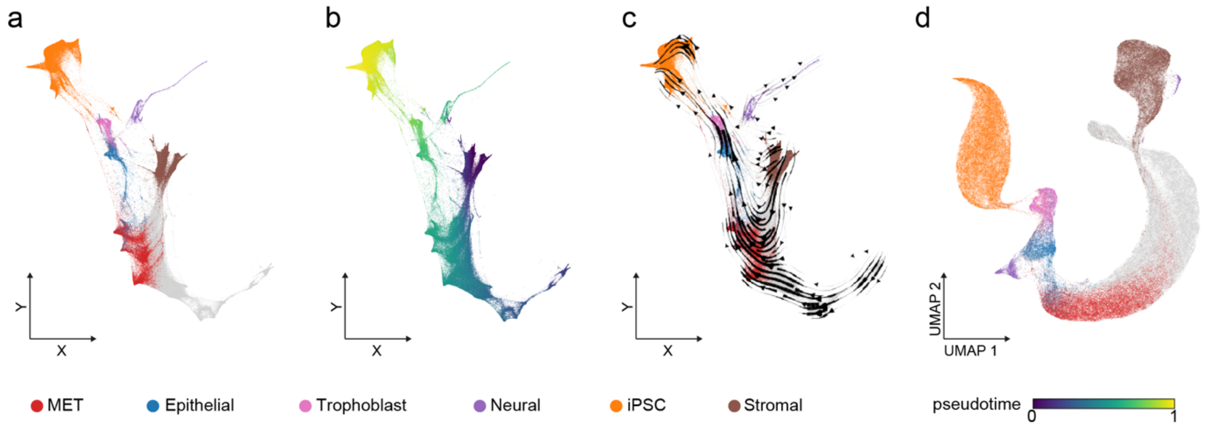


**Supplementary Figure 7 | scTour captures the cellular dynamics during iPSC reprogramming in mice.**

**a,** Force-directed layout embedding (FLE) visualizing cell states from 251,203 cells collected during iPSC reprogramming. The FLE coordinates in **a-c** and cell annotations are from the original study. MET, mesenchymal-to-epithelial transition.

**b-d,** FLE (**b-c**) and UMAP (**d**) visualizations of the inferred developmental pseudotime (**b**), transcriptomic vector field (**c**), and latent space (**d**) from the scTour model trained using 20% of all cells.

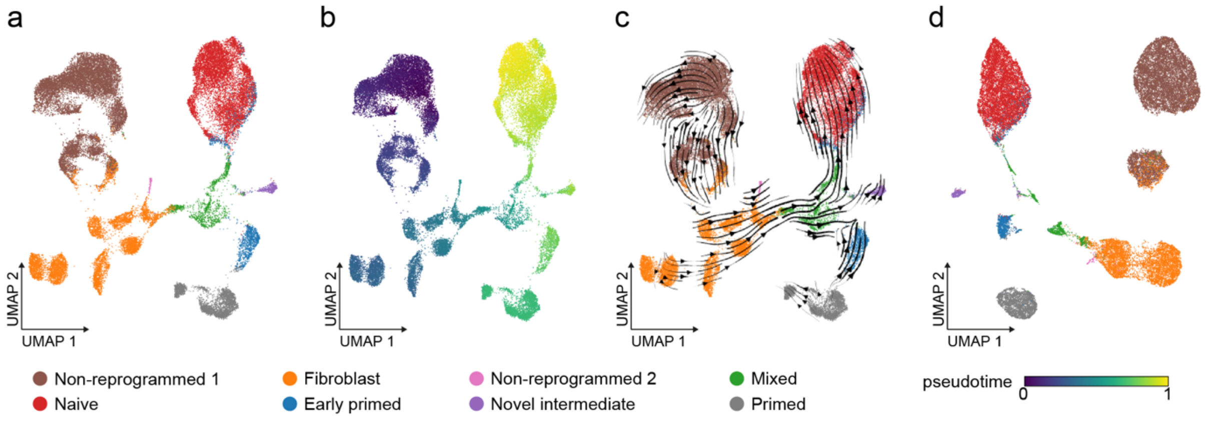


**Supplementary Figure 8 | scTour captures the cellular dynamics during iPSC reprogramming in humans.**

**a,** UMAP visualization of cell states from 36,597 nuclei collected during iPSC reprogramming by snRNA-seq. The UMAP coordinates in **a-c** and cell annotations are from the original study.

**b-d,** UMAP visualizations of the inferred developmental pseudotime (**b**), transcriptomic vector field (**c**), and latent space (**d**) from the scTour model trained using 60% of all cells.

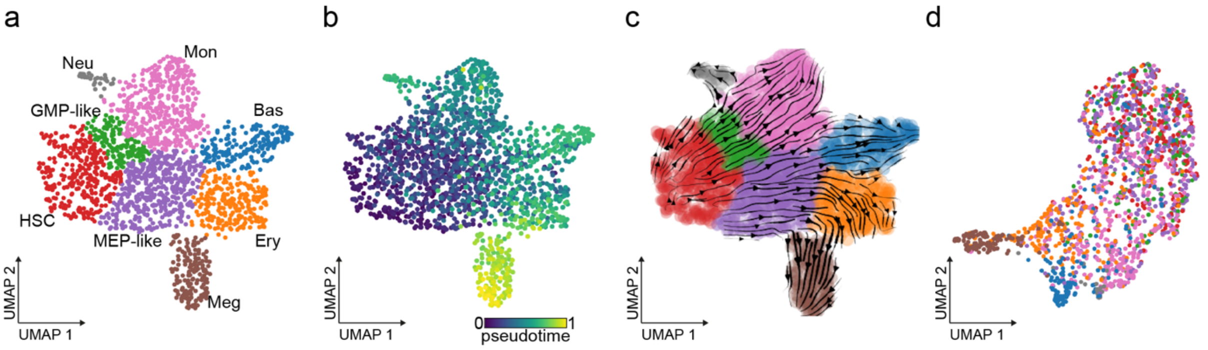


**Supplementary Figure 9 | scTour captures the developmental cellular dynamics during hematopoiesis.**

**a,** UMAP visualization of cell lineages from 1,947 cells profiled by scNT-seq during hematopoiesis. The UMAP coordinates in **a-c** and cell type annotations are from the original study. HSC, hematopoietic stem cell; Meg, megakaryocyte; Ery, erythrocyte; Bas, basophil; Mon, monocyte; Neu, neutrophil; GMP-like, granulocyte and monocyte progenitor-like cells; MEP-like, Meg and Ery progenitor-like cells.

**b-d,** UMAP visualizations of the inferred developmental pseudotime (**b**), transcriptomic vector field (**c**), and latent space (**d**) from the scTour model trained using 90% of all cells.

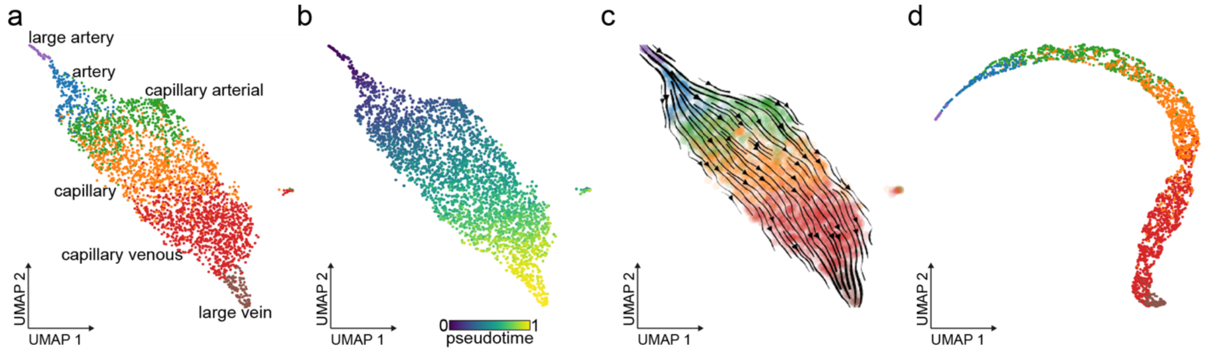


**Supplementary Figure 10 | scTour captures the anatomical topography of brain endothelial cells.**

**a,** PCA space-based UMAP visualization of cell clusters from 3,105 endothelial cells collected from the mouse brain. The cell type annotations are from the original study.

**b-d,** UMAP visualizations of the inferred developmental pseudotime (**b**), transcriptomic vector field (**c**), and latent space (**d**) from the scTour model trained using 90% of all cells.

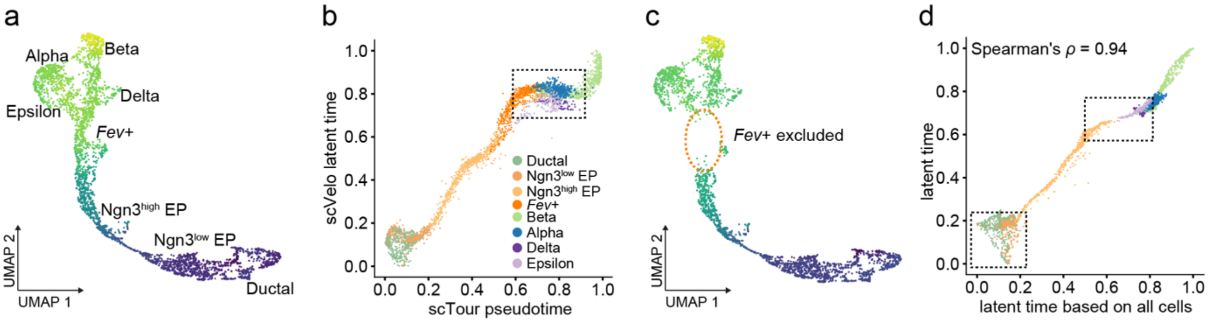


**Supplementary Figure 11 | Superiority of scTour's pseudotime over scVelo's latent time in characterizing a discontinued process.**

**a,** UMAP showing the latent time inferred by scVelo for all the cells along the endocrinogenesis process.

**b,** Comparison of scTour's pseudotime (x axis) with scVelo's latent time (y axis). Dots represent cells coloured by their identities of cell types. The rectangle marks the continuous differentiation process from *Fev*+ endocrine cells to terminal cell fates captured by scTour's pseudotime but not by scVelo's latent time.

**c,** UMAP showing the latent time inferred by scVelo when *Fev*+ endocrine cells are excluded.

**d,** Comparison of the latent time from **c** (inferred from cell subset with *Fev*+ endocrine cells held out, y axis) with that from **a** (inferred from all cells, x axis). The bottom-left rectangle shows the early progenitor progression which is not properly delineated by scVelo's latent time when *Fev*+ endocrine cells are held out. The upper rectangle marks the inability of scVelo to capture the transcriptomic discontinuity between Ngn3$^{high}$ EPs and terminally differentiated cells when the intermediate *Fev*+ endocrine cells are excluded.

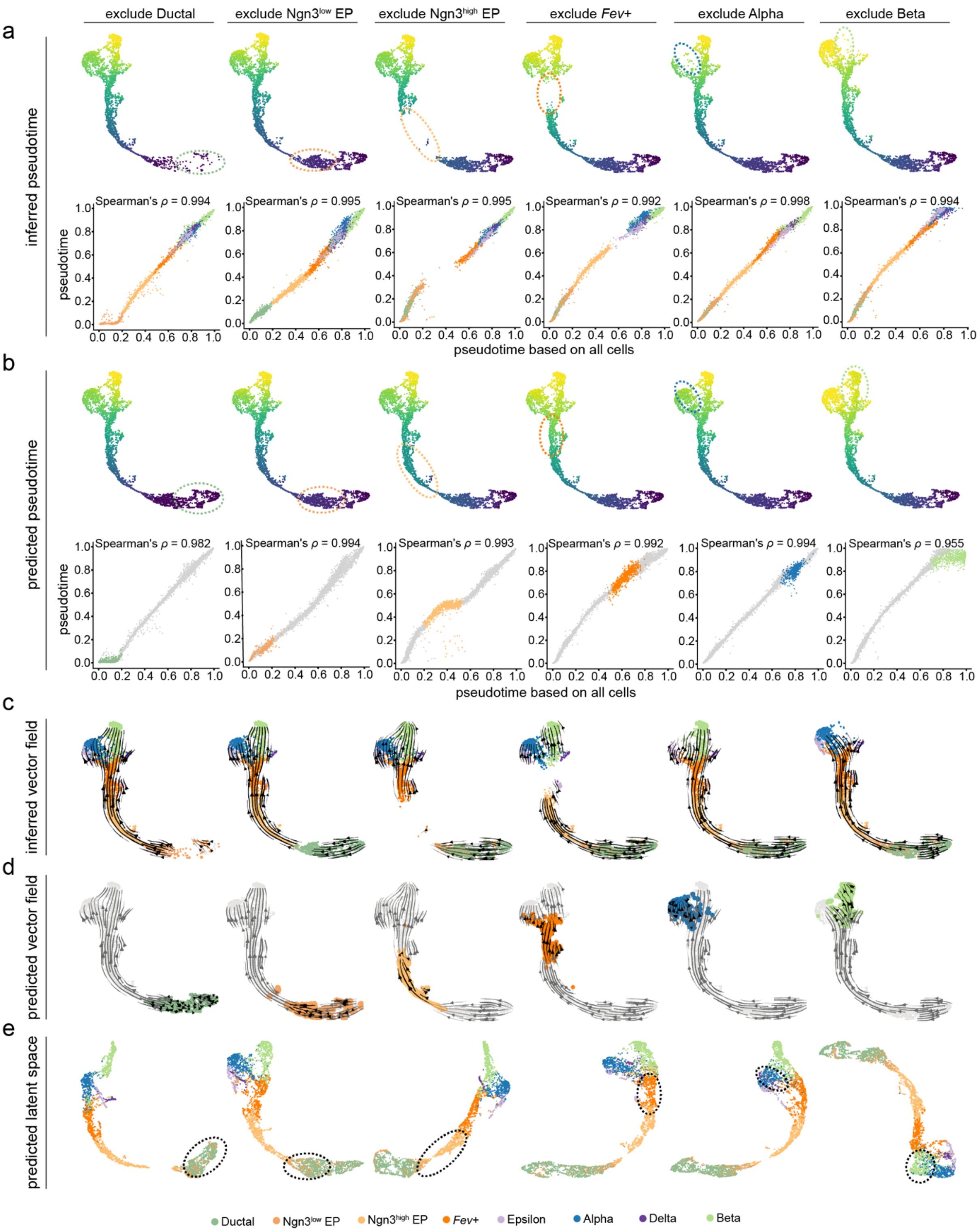


**Supplementary Figure 12 | scTour predicts the cellular dynamics of unseen cellular states regardless of their positions along the developmental process.**

**a,** UMAP visualizations of the developmental pseudotime inferred by the scTour model trained with a certain cellular state held out (upper panels). Scatter plots in the bottom panels show the comparisons with the pseudotime estimates from the model trained with all cells. The

Spearman correlation coefficients calculated between the two sets of estimates are shown on top.

**b,** Upper panel: UMAP visualizations showing the predicted pseudotime for the held-out cellular states (dotted circles). Lower panel: scatter plot showing the comparisons of the prediction with the ground truth. The Spearman correlation coefficients calculated between pseudotime in x and y axes are shown on top.

**c,** UMAP representations of the transcriptomic vector fields inferred by the scTour model trained with a certain cellular state held out.

**d,** UMAP representations of the predicted transcriptomic vector fields for the held-out cellular states.

**e,** UMAP visualizations based on the predicted latent representations for the held-out cellular states (dotted circles) and those inferred from the training cells.

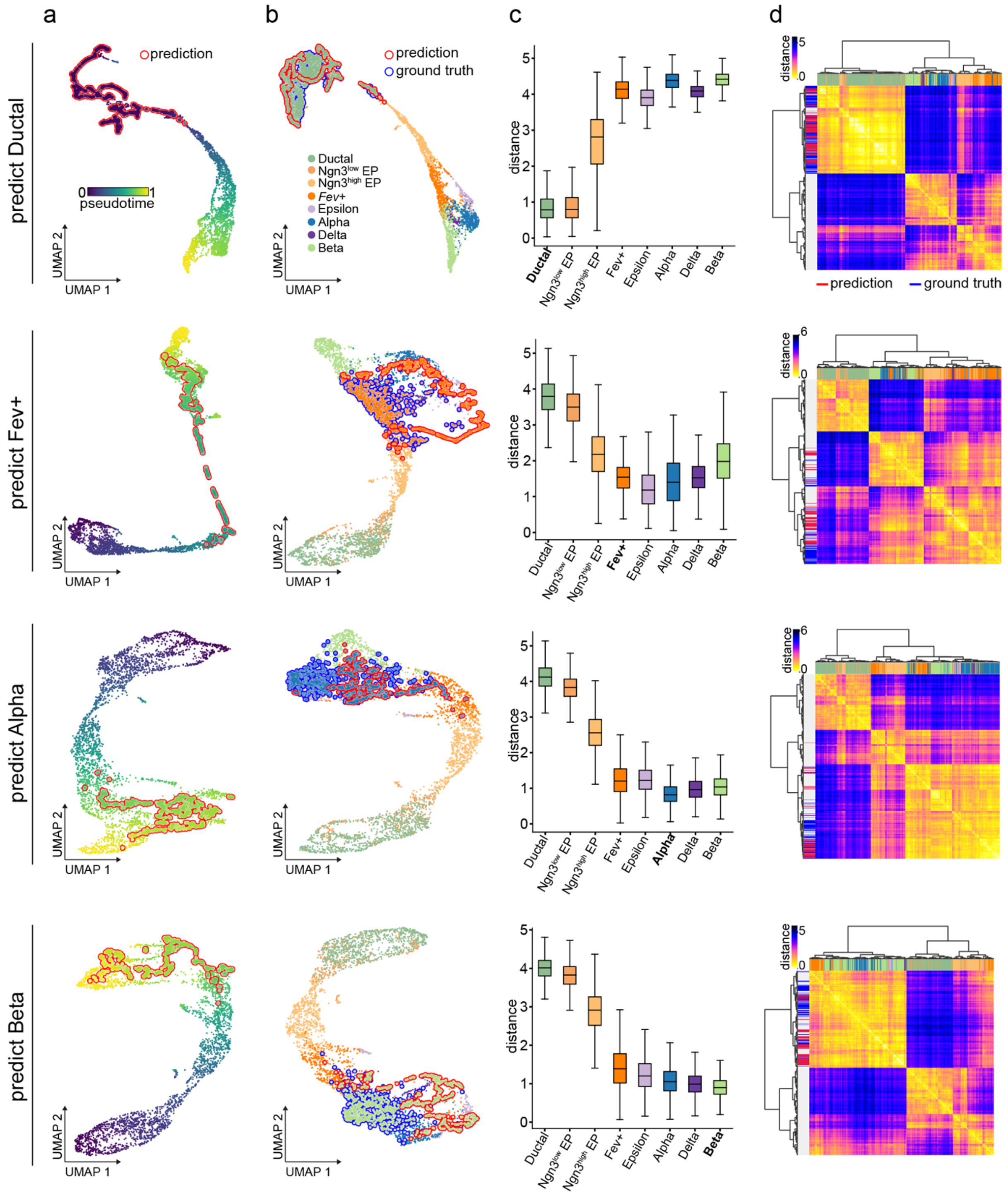


**Supplementary Figure 13 | scTour reconstructs the transcriptomic space at different developmental stages**

**a,** UMAP visualizations of the reconstructed latent representations for the held-out cellular states given their expected developmental pseudotime (red outline) and the latent space inferred from the training cells.

**b,** UMAP visualizations based on the latent representations from the reconstructed cells (red outline), ground-truth cells (blue outline), and remaining cells. Cells are colour-coded by cell types.

**c,** Box plots showing the Euclidean distances calculated between the reconstructed latent representations of the held-out cellular state and those from the ground truth and remaining cellular states, with the medians, interquantile ranges, and 5th, 95th percentiles indicated by centre lines, hinges, and whiskers, respectively.

**d,** Unsupervised hierarchical clustering of the reconstructed cells along with all the other cells based on their Euclidean distances in the scTour latent space. Column colours of the heatmap mark the cell types and row colours denote the reconstructed (red), ground-truth (blue), and remaining (light grey) cells. The colour shades of the heatmap indicates the Euclidean distance.

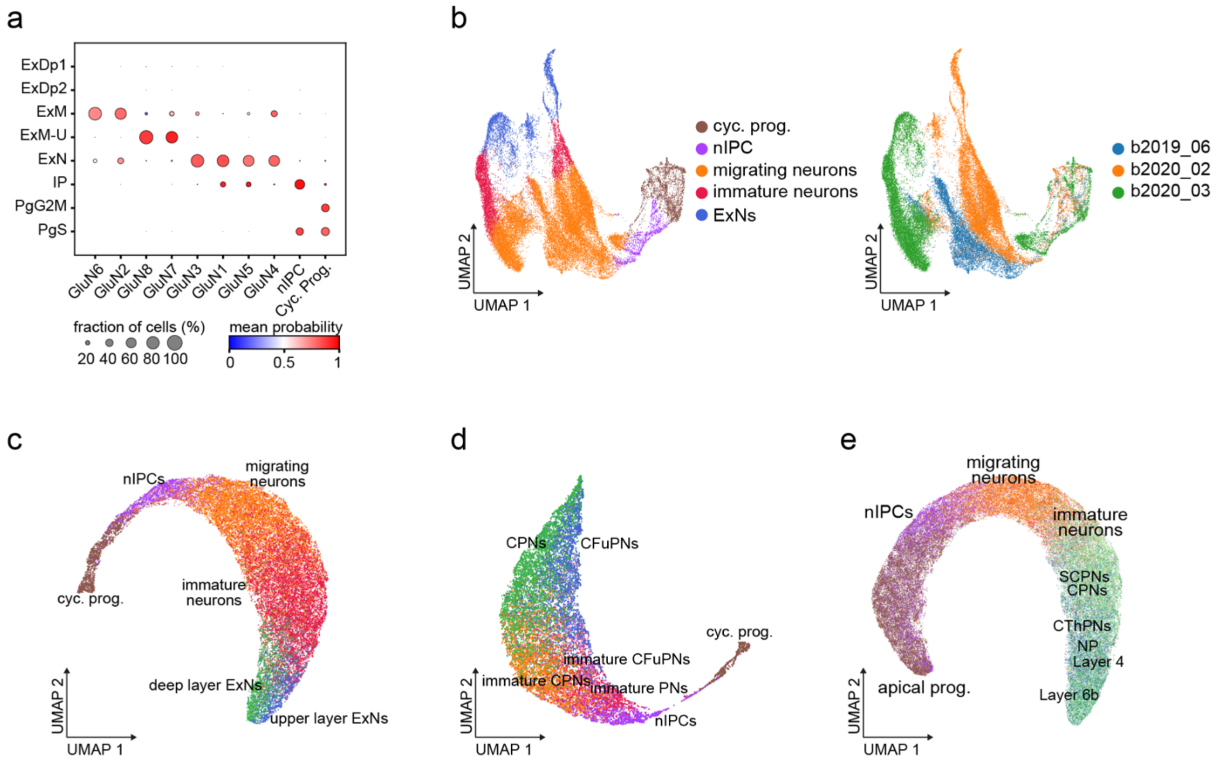


**Supplementary Figure 14 | scTour predicts the latent space of unseen datasets.**

**a,** Dot plot showing the cell type label transfer from Polioudakis et al. (row) to cells from Trevino et al. (column). Across rows of each column, dot size indicates the proportion of cells assigned to a given cell type from Polioudakis et al. and colour represents the average probabilities calculated by CellTypist.

**b**, PCA space-based UMAP visualizations of the cell types from the training dataset, with cells coloured by cell types (left) and sample batches (right).

**c-e,** UMAP visualizations of the predicted latent representations for the three test datasets: the developing human cortex (**c**), the human brain organoid (**d**), and the developing mouse cortex (**e**). cyc. prog., cycling progenitors; nIPCs, neuronal intermediate progenitor cells; ExNs, excitatory neurons; PNs, projection neurons; CPNs, callosal projection neurons; CFuPNs, corticofugal projection neurons; CThPNs, corticothalamic projection neurons; NP, near projecting; SCPNs, subcerebral projection neurons; apical prog., apical progenitors.

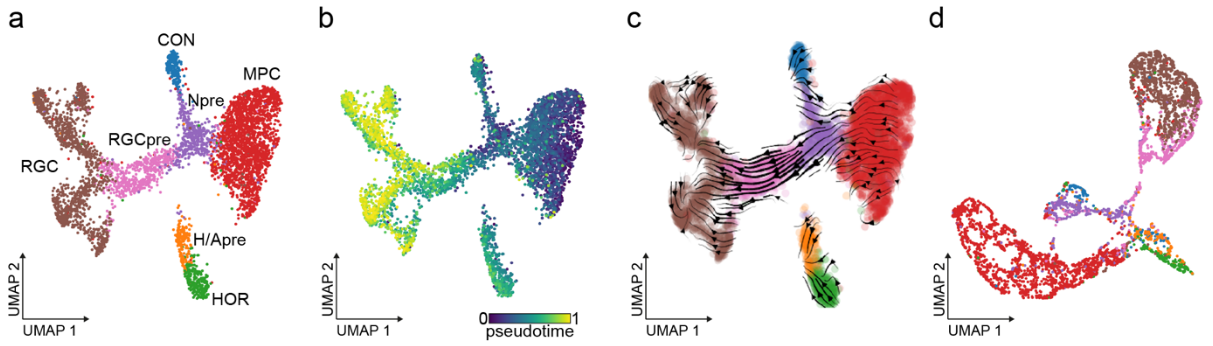


**Supplementary Figure 15 | Application of scTour to scATAC-seq data.**

**a,** Latent semantic indexing (LSI)-based UMAP visualization of the cell types identified from 4,883 cells of the human fetal retina measured by scATAC-seq. RGC, retinal ganglion cells; CON, cones; NPre, neurogenic precursors; RGCpre, RGC precursors; H/Apre, the horizontal/amacrine precursors; MPC, multipotent progenitor cells; HOR, horizontal cells.

**b-d,** UMAP visualizations of the inferred developmental pseudotime (**b**), epigenetic vector field (**c**), and latent space (**d**) from the scTour model trained using 90% of cells and top 25% of peaks (34,670 genomic regions).
